# Automated Purification of DNA Origami with SPRI Beads

**DOI:** 10.1101/2023.07.05.544573

**Authors:** Chalmers Chau, Gayathri Mohanan, Iain Macaulay, Paolo Actis, Christoph Wälti

## Abstract

DNA origami synthesis is a well-established technique and has been employed in various applications. The synthesised origami must be purified to eliminate the excess materials such as DNA oligos and other molecules. While several purification techniques are routinely used, they all have limitations, and none can be automated to simultaneously handle large numbers and quantities of samples. Here we introduce the use of solid-phase immobilisation (SPRI) beads as an easy-to-adopt, scalable, high-throughput and automation-compatible method to purify DNA origami. Not only can this method remove excess oligos and biomolecules with comparable yield to existing methods while maintaining high structural integrity of the origami, but it also allows an automated workflow to simultaneously purify large numbers of samples within a limited time. We envision that the SPRI beads purification approach will improve the scalability of DNA nanostructures synthesis both for research and commercial applications.

## Introduction

The use of DNA as a building block for the creation of nanoscale materials is the foundation of the field of DNA nanotechnology^[1]^. For example, the DNA origami technique involves the combination of a long ssDNA scaffold with hundreds of short oligonucleotides, “staple” strands, via Watson-Crick base pairing to assemble rationally designed nanostructures^[1b]^. It has found numerous applications in biophysical research, clinical diagnostics, and in cell biology^[1d, 2]^. Additionally, the programmability of DNA enables the precise modification of the origami nanostructures with a range of functional biomolecules ^[1d, 3]^. While the assembly and programmability is well-established in the preparation of such DNA nanostructures, the purification of the desired products from the excess materials used during the assembly, such as staples and functional molecules^[1c]^, is still challenging.

A wide range of purification techniques have been developed and are used routinely. These methods include gel extraction, poly(ethylene) glycol (PEG) precipitation, molecular weight cut-off (MWCO) membrane filtration and spin column-based filtration method ^[1c, 1d, 4]^. Agarose gel electrophoresis separates the slow-migrating folded DNA nanostructure as a distinct band from the faster migrating staples. The desired band can be excised from the gel and the product extracted ^[4a]^. An alternative approach relies on the property of polyethylene glycol (PEG) to induce DNA precipitation ^[4b]^. The third and fourth widely used method rely on filtration using MWCO membrane and chromatography resins, respectively ^[1c]^.

Most of these methods are not suitable for high volume purifications and often require manual operations precluding their automation with liquid handling robots ^[1c]^. This is a major obstacle for the scaling up of their production and implementation in industrial settings. Here, we report the use of solid-phase reversible immobilisation beads (SPRI) ^[5]^ for the manual as well as automated purification of a range of DNA origami at high concentration from excess staples and proteins (Figure 1, Figure S1). The SPRI beads are paramagnetic microparticles modified with carboxyl groups which can reversibly bind to DNA and are widely employed in transcriptomics for DNA size selection and purification prior to sequencing ^[5-6]^. Compared to other methods, this technique does not require centrifugation ^[1c, 4b, 7]^ or chemical modifications of the origamis ^[8]^, which present limitations for automated large-scale implementations, and we demonstrate the readiness of this method for scaling up by performing an automated purification of 96 DNA origami samples with a liquid handling robot ^[9]^. We envisage that the SPRI clean-up method will further enable the commercial exploitation of DNA nanostructures by allowing their high throughput purification.

**Figure 1.**
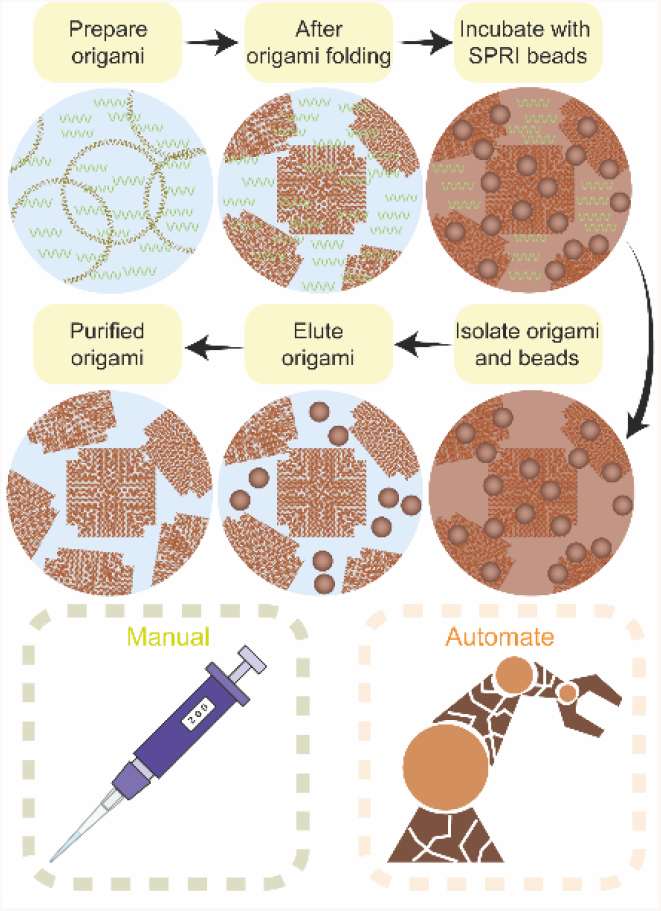
Schematic illustration of SPRI clean-up of DNA origami. The DNA origamis are mixed with the SPRI beads at a specific beads-to-volume ratio, followed by their separation with an external magnet. The origami beads pellet is then washed with ethanol followed by elution in the DNA origami storage buffer.

## Results and discussion

SPRI clean-up is an effective method to purify DNA nanostructures from excess staples. An 88 nm × 88 nm 4-fold symmetrical tile (4FST) DNA origami ^[10]^ was selected and used throughout this study as the model DNA nanostructure for the validation of the SPRI clean-up. To achieve high quality purification, it is important to optimize the volume ratio between sample and SPRI beads ^[6b, 11]^. The volume ratio was investigated between 0.4X and 4.0X to identify the optimal in terms of efficient removal of excess staples following the DNA origami assembly. We performed agarose gel electrophoresis to identify the optimal volume ratio by observing whether staples could still be detected in the gel (Figure 2A). The folded 4FST origami band migrated slower compared to the scaffold due to differences in its mobility. The densitometric lane profile of all volume ratios was analysed (Figure 2B, Figure S5A), and the integrity of the origamis for all ratios were checked with Atomic Force Microscopy (AFM) imaging (Figure 2C, Figures S6-S12) demonstrating that the SPRI beads did not affect the integrity of the origamis. We confirmed from agarose gel electrophoresis (Figure 2A, B) that volume ratios higher than 1X caused incomplete purification as is demonstrated by the presence of excess staples (lower molecular weight materials) in the sample after purification. Ratios of 1X or lower generated the purest origamis and no staples were observed in the gel electropherogram, with a total mass of origami ranging from ∼700 ng to ∼1200 ng in a 40 μl reaction (Figure S5B). We chose the 0.8X ratio for all downstream testing and successfully purified three other origami designs (dimer 4FST, 4FSF, and frame) to demonstrate that the method is universally applicable (Figures S13-16).

**Figure 2.**
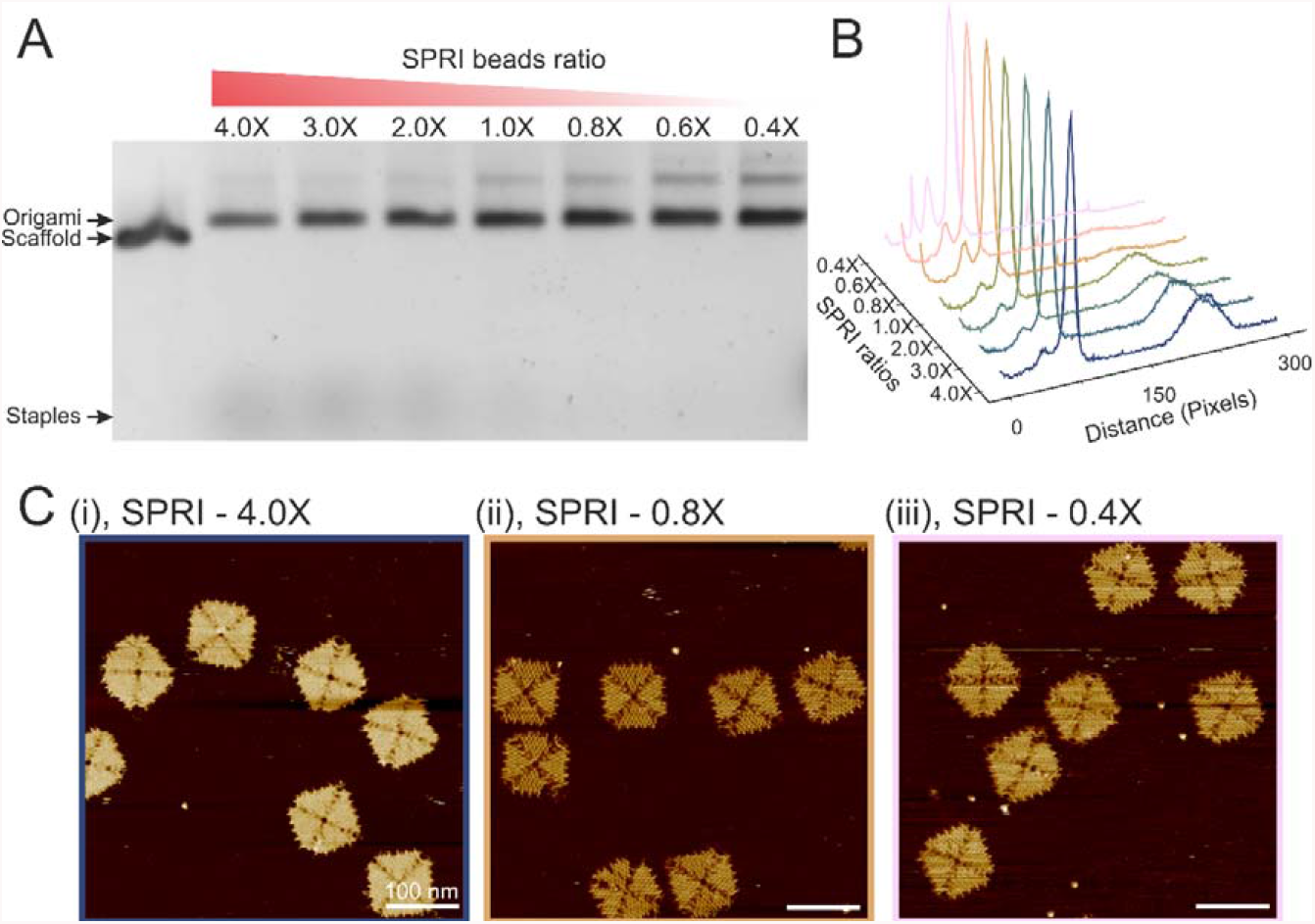
Purification of DNA origami using different ratio of SPRI beads to origami. **A**, Agarose gel of the 4FST origamis after SPRI purification at different volume ratio. **B**, cross-sectional lane profile plotted from A. **C**, AFM images of (i), 4X (ii), 0.8X (iii), 0.4X ratio purified 4FST origami (scale bar: 100 nm).

While the purification yield was excellent, we observed that the SPRI beads-purified origami clumped together into a cluster (Figure S3). We hypothesised that the formation of clumps was a result of the dehydration caused by the ethanol wash during the purification, as previous studies reported that high percentages of alcohol led to the condensation and precipitation of DNA due to electrostatic interactions ^[12]^. A thermal de-clumping step was introduced, this step allowed the origami clump to be removed successfully as evidence in the agarose gel and AFM images (Figures S3). In addition to the SRPI beads used here, an additional commercially available SPRI beads were also tested, and both successfully purified origami (Figure S4).

Various techniques have been used to purify DNA origami and we performed a systematic study to benchmark the SPRI beads purification approach against several established purification methods. These methods include the S-400 HR spin column filtration, two different 100 kDa MWCO filtrations, PEG precipitation, phase separation, ethanol precipitation and size exclusion chromatography (SEC) ^[1c, 4b, 7, 13]^. The uncleaned origami samples and origami purified using each method were analysed via agarose gel electrophoresis (Figure 3A). The uncleaned samples showed a band corresponding to low molecular weight products that were attributed to the excess staples. All methods, except 100 kDa MWCO-2, phase separation and ethanol precipitation, had successfully purified the DNA origami from the unreacted staples as based on gel electropherogram and its associated densitometry lane profile (Figure 3A, B). We observed clumping in the gel well with the ethanol precipitation method, which was similar to what we had observed in SPRI clean-up prior to the thermal de-clumping (Figure S3). Baptist *et al*. recently observed similar aggregation after purifying origamis via PEG precipitation, which they attributed to stacking interactions between origamis after prolonged centrifugation ^[14]^. Although no centrifugation was involved with our SPRI clean-up method, stacking interactions could also have been enhanced during the ethanol wash step in our method, leading to the aggregation of origamis.

**Figure 3.**
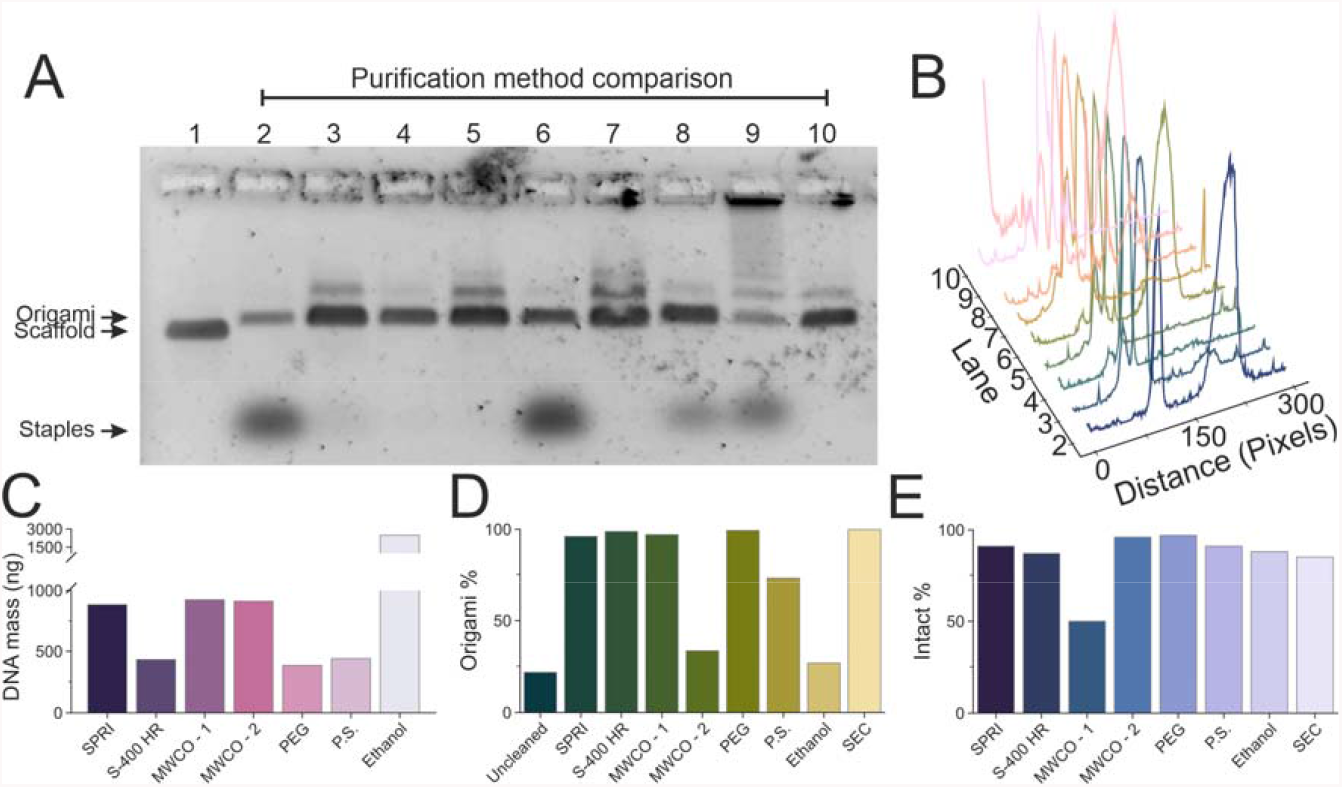
Comparison of 4FST origami purification using different methods. **A**, Agarose gel image of origami purified using different methods. Lane labels: 1, M13mp18 scaffold; 2, uncleaned 4FST tile; 3, 0.8X SPRI ratio; 4, S-400 HR spin column; 5, 100 kDa MWCO-1 filter; 6, 100 kDa MWCO-2 filter; 7, PEG precipitation; 8, Phase separation; 9, Ethanol precipitation; 10, SEC column purified. **B**, Densiometric lane profile plotted from A. **C**, Purification yield calculated from absorption spectroscopy. **D**, Percentage of DNA origami calculated from B. **E**, Percentage of intact origami after purification using each method quantified from AFM imaging.

When the total DNA origami yield was measured (Figure 3C), the ethanol precipitation method delivered the highest yield followed by the two MWCO methods, but a significant number of staples were still present in the sample following purification (Figure 3A). The SPRI clean-up yielded similar amount of purified DNA origami as MWCO-1 and MWCO-2. When we quantified the percentage of folded DNA origami to staples based on gel densitometry, we found that SPRI beads had a comparable performance against other methods including the S-400 HR column, MWCO-1, PEG precipitation and SEC (Figure 3D), indicating that all these methods were highly effective in purifying DNA origamis from unreacted staples. Lastly, we used AFM imaging to assess the structural integrity of the origamis following purification for each of the methods discussed above (Figures S17-23). All purifications retained 80% and above origami structural integrity apart from MWCO-1 at 50% (Figure 3E), where significant deformation of the 4FST tile structure could be observed (Figure S18).

In summary, we conclude that the SPRI purification method is a comparable technique in terms of yield, DNA origami purity, and structural integrity to the best performing purification methods.

There are a wide range of applications that require the functionalisation of DNA origami with materials such as fluorophores, proteins, and nanoparticles ^[1d, 3, 15]^. To enable high functionalisation yields, the excess material needs to be purified to prevent interference with downstream applications, while retaining the functionalised DNA origami ^[16]^. SPRI clean-up is not only an effective method to purify DNA nanostructures from excess staples but can also be employed to remove proteins. We used the C-reactive protein (CRP) as a model protein as it had a considerable size (125 kDa), and such proteins are difficult to be purified using conventional methods such as membrane purification ^[1d, 16]^. CRP was added to the SPRI purified 4FST origami in 2:1 ratio, and this was then cleaned by 0.8X ratio of SPRI beads (Figure 4A). Multiple rounds of cleaning were carried out with the origami-CRP mixture to ensure the complete removal of CRP. Surprisingly, we observed that CRP was completely removed during the first cleaning step as could be seen by SDS-PAGE gel, where the band corresponding to CRP had disappeared (Figure 4B). AFM images also confirmed the removal of CRP from the background and predominantly intact origamis were observed after the clean-up (Figure 4C, Figure S24). Furthermore, we assessed the performance of the SPRI purification to retain intact origami from multiple rounds of purification (Figure 4D, Figure S25-27) at different SPRI beads ratio and using S-400 HR spin column filtration for comparison. We found that the SPRI bead purification retained origami structures even after multiple rounds of clean-up with ∼500 ng left after three rounds, in contrast to ∼200 ng left after S-400 HR spin column filtration. These findings demonstrate that SPRI bead clean-up is an effective method to purify DNA origami from other excess materials.

**Figure 4.**
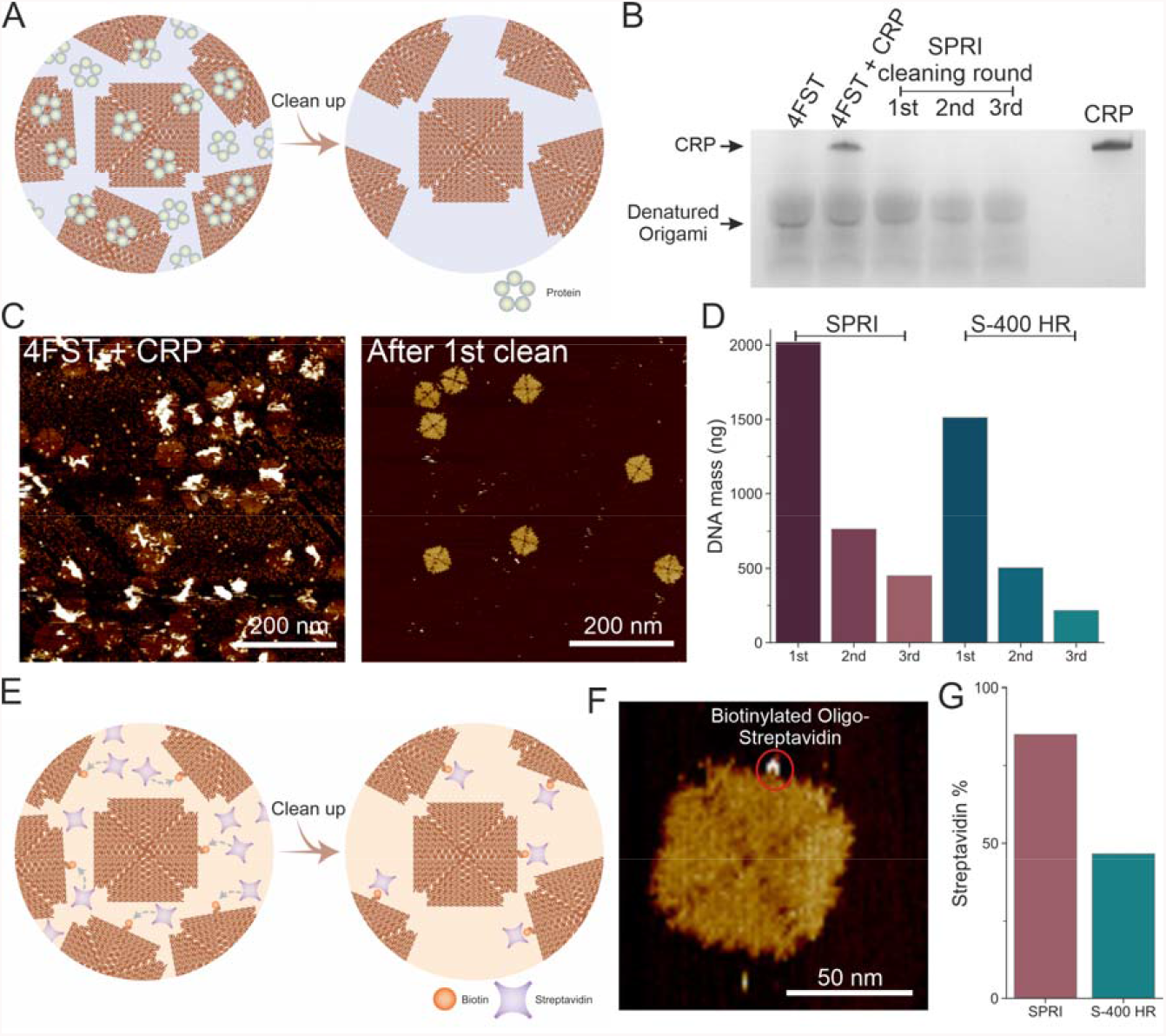
SPRI assisted removal of excess proteins. **A**, Schematic illustration of removing excess proteins from origami. **B**, Silver stained SDS-PAGE gel of 1^st^, 2^nd^ and 3^rd^ round cleaned and uncleaned origami, CRP as loading control. **C**, AFM images of origamis with CRP before and after 1^st^ round of SPRI cleaning (scale bar: 200 nm). **D**, Yield by DNA mass of origamis after three rounds of cleaning using 0.8X SPRI beads and S-400 spin columns. **E**, Schematic illustration of streptavidin conjugation to origami containing biotinylated staples and the subsequent removal of excess streptavidin using 0.8X SPRI beads. **F**, AFM image of streptavidin conjugated to a biotinylated 4FST tile. **G**, bar graph indicating the total yield of streptavidin functionalised 4FST origami purified using 0.8X ratio SPRI beads and S-400 spin columns.

In scenario where excess materials were used to functionalise the DNA origami, it’s important that the purification method does not interfere or reverse with the functionalisation. To test this, we investigated the functionalization of a DNA origami with streptavidin through interaction with a biotinylated staple strand as a model system ^[1d, 3]^ (Figure 4E, F). We employed the SPRI beads clean-up procedure alongside the S-400 HR spin column filtration on the biotinylated origami after incubating the biotinylated origami with streptavidin at 2:1 molar ratio. The percentage of the streptavidin retained on the origami was quantified by AFM and we found that SPRI purification led to a high percentage (∼85%) of streptavidin being retained on the origami compared to S-400 HR filtration where more than half of the origami had streptavidin bound (Figure 4G, Figure S28), demonstrating the advantage of SPRI beads for origami functionalisation clean-up.

The key advantage of SPRI beads-based purification is that it is compatible with large-scale robotic purification. Here, we demonstrate that the SPRI clean-up can be automated by implementing the protocol on an automated liquid handling robot, which enables the purification of DNA origami from 96 reactions simultaneously within 30 minutes (Figure 5A). Following the folding reaction inside a 96 wells PCR plate, the liquid handling robot was programmed to perform the SPRI clean-up procedure, and after the elution of the origami, the plate was transferred to thermocycler for the de-clumping step. Afterwards, we analysed the purified products via agarose gels (Figure 5B, Figures S29, 30) and AFM imaging (Figure 5C, Figure S31). This clearly demonstrates that the automation routine performs as well as the manual pipetting.

**Figure 5.**
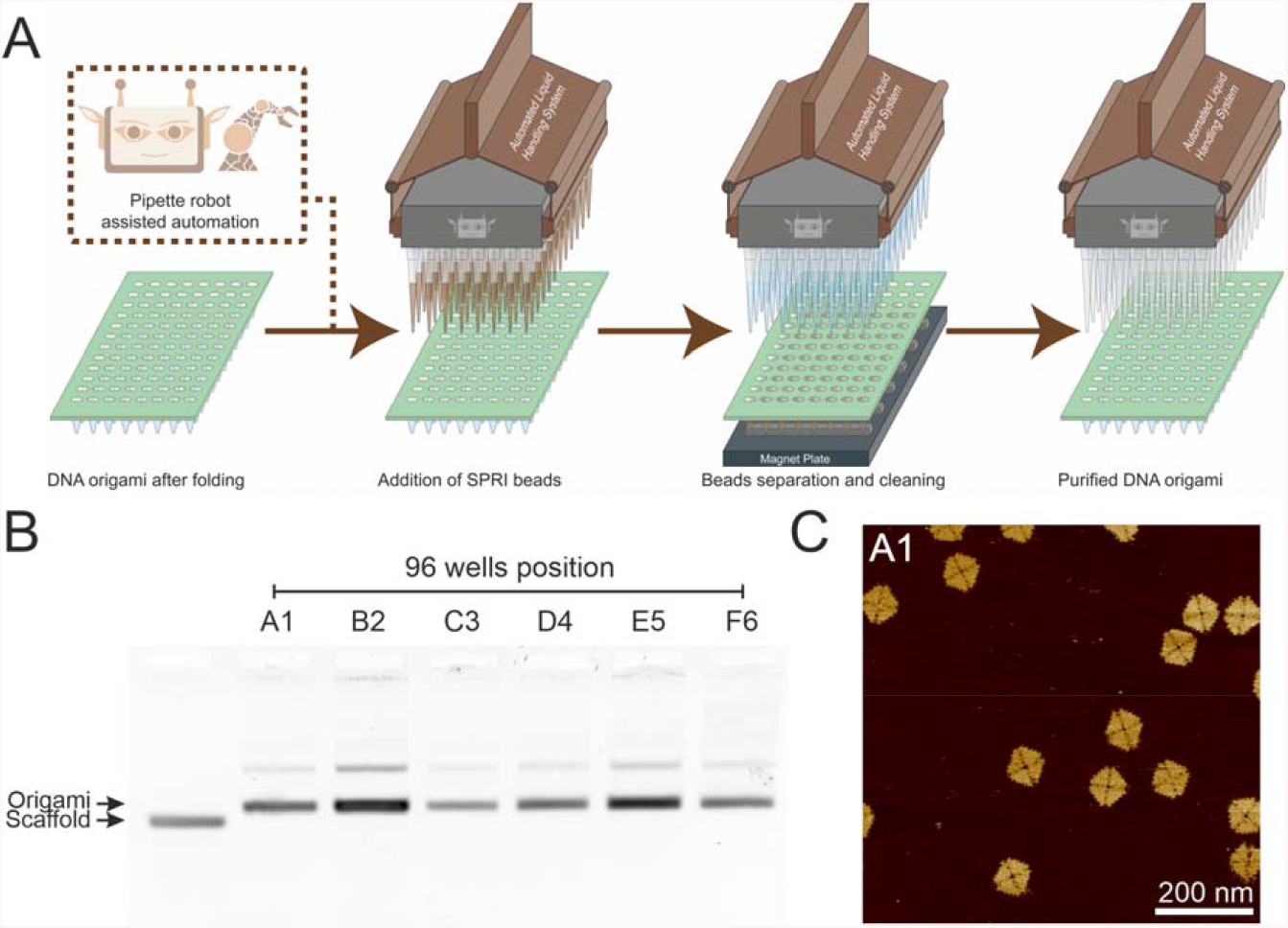
Automated preparation of purified 4FST origami. **A, Schematic illustration of** the liquid handling robot **performing** the **SPRI purification** procedure on the origami samples. **B**, Random 6 samples of the origami purified via automation. **C**, AFM image of sample taken from well A1 (scale bar: 200 nm).

The SPRI beads-based purification procedure requires only basic instruments, reaction tubes, pipettes and magnets ^[5-6]^, thus this procedure is automation compatible where minimal manual interference is required ^[9]^. In contrast, methods that are based on precipitation or filtration rely on centrifugation ^[1c, 4b, 7]^ which is the main obstacle in automation. Upgrading to robotic cleaning can aid in automating repetitive cleaning steps thus making it time efficient, less error prone and providing increased throughput. Automation helps in improved scalability as it can handle large quantities of samples, additionally, lyophilisation can be implemented after the SPRI purification to store and transport purified DNA nanostructures at large quantities ^[14, 17]^, which can further enable the commercial exploitation of DNA nanostructures.

Magnetic beads have been used before to purify origami, but this approach relies on the chemical coupling of the DNA origami onto the surface of the magnetic beads ^[8, 13]^. The method established in this work is universal and does not require chemical coupling of the DNA origami onto the beads. It can be utilised for purifying origami structures of any design and functionalised origami after their synthesis and conjugation irrespective of the oligo or functional molecule used. Previous studies have suggested that the buffer component of the SPRI beads could be further refined to increase the size-selectivity of the beads ^[11]^, the composition of the SPRI mixture can be further modified in the future to further optimise the clean-up efficiency.

## Conclusion

We have demonstrated the use of SPRI beads, which are well-established in sequencing, as an innovative technique to purify DNA origami. We analysed the efficiency of this technique at different ratio of beads to origami and selected the optimum ratio for purification. This method is universal and can be applied for wide range of origami designs. We then compared the SPRI technique with existing purification methods, and it can be used as a reliable method with comparable yields to the available methods. Moreover, we anticipate the use of this method to remove excess materials such as proteins and retaining high proportion of materials functionalised origamis. Lastly, we advocate the possibility of expanding origami purification from bench level to industrial level by automating the SPRI clean-up procedure. Successful implementation of high-throughput automation to prepare purified origami of various designs means increased scalability and adaptability not only for research, but also for the industry sector, assisting the development of DNA nanotechnology.

## Methods

For method details, please refer to the supporting information for details.

## Data availability

Data (AFM images, DNA origami design file and staples list, images used for the calculation of the origami intact percentage) supporting this work can be freely accessed via the University of Leeds data repository: https://doi.org/10.5518/1369

## Supporting Information

Supporting information include detailed methods for the folding of DNA origami, purification of DNA origamis with different methods, AFM, automation set-up and supporting figures.

## Author Contributions

C.C. and G.M. contributed equally at this work. C.C. designed the research study, G.M. performed the experiments. C.C. and G.M. analysed data. C.C. illustrated all schematics. I.M. helped with the robot automation. P.A. and C.W. helped with data analysis, supervised the work and acquired the funding. All authors wrote and corrected the manuscript.

## Acknowledgement

We thank the group of bioelectronics member for providing insightful feedbacks. We also want to thank for all the automation technical supports received from Earlham Institute’s people.

## Abbreviation

SPRI: Solid Phase Reversible Immobilization
PEG: poly(ethylene) glycol
MWCO: molecular weight cut-off
AFM: Atomic Force Microscopy
SEC: size exclusion chromatography
CRP: C-reactive protein

