## Supporting Information for "Automated Purification of DNA Origami with SPRI Beads"

### Section 1: Supporting Methods

All chemicals were acquired through Sigma-Aldrich at ACS grade unless otherwise stated. All concentrations stated here are at final concentration unless otherwise stated. All buffers (except buffers containing PEG or Dextran) were sterile filtered with either 0.1μm (16553K; Sartorius) or 0.22μm (SLGP033RS; Millipore) syringe filter.

#### DNA origami production

All DNA strands used throughout this study were ordered from Integrated DNA Technologies (IDTDNA) with standard desalting service. The caDNAno (2.4.7) software was used to design the DNA origami structures [1]. All origamis were folded using the 7249 nt M13mp18 circular single-stranded DNA scaffold (N4040S; NEB). All folded origamis were stored at 4°C or immediately purified.

The caDNAno files of the origami designs, exported staple lists and the M13mp18 sequence used to export the staples were included in the repository associated with this study.

##### 4-Fold Symmetrical Tile (4FST) and the biotinylated 4FST origami

The design of the 4-Fold Symmetrical Tile (4FST) DNA origami used here follows the design published by Tikhomirov *et al.* [2]. The 4FST DNA origami staples are categorized into 4 staple mixtures: interior, bridge, edge and negation, each mixture is a mixture of all relevant staples.

To fold 40μl of the 4FST DNA origami, the folding mixture contained: 10nM M13mp18 scaffold, 12.5mM magnesium acetate, 75nM bridge mixture, 75nM bridge mixture, 75nM edge mixture, and diluted in 10mM tris-HCl, 1mM Ethylenediaminetetraacetic acid (EDTA) at pH 8.0 (1X TE buffer) (93283; Sigma-Aldrich). The folding of the 4FST DNA origami was carried out with a thermal cycler by heating the folding mixture to 90°C for 2 minutes, followed by a gradual temperature decrease from 90°C to 20°C at the rate of 1°C per minute. The temperature was then further decreased down to 4°C until use.

For the biotinylated 4FST DNA origami, one of the standard edge staples (Edg-T1R10C7-DHP) in the edge staple mixture was replaced with a 5' biotinylated version of the same staple. To fold the biotinylated 4FST, the same procedure as the standard 4FST was used. After the purification of the biotinylated 4FST origami, a total of 20nM streptavidin (434301; Thermo Fisher) was added to the 4FST origami and incubated for 10 minutes at room temperature, then the biotinylated 4FST and streptavidin mixture was subjected to purification again.

##### Dimeric 4-Fold Symmetrical Tile (4FST)

To generate the dimer 4FST, the edge was replaced with a different set of edge staples that allows the hybridisation between two 4FST [2]. After the folding of the two separate 4FST origamis, the two samples were mixed at 1:1 volume (including all excess staples). The mixture was then heated to 55°C and the temperature was then gradually decreased to 20°C at a 1°C/min rate. The temperature was then further decreased down to 4°C until use.

##### 4-Fold Symmetrical Frame (4FSF) origami

The basic design principle of the 4-Fold Symmetrical Frame follows the 4FST, thus the folding mixture components and the steps were the same as 4FST except for the differences in staple sequences.

##### Frame origami

The design of the frame origami used here followed previously published design by Raveendran *et al.* [3]. All the staples were pooled together into the staple mixture. The origami folding mixture contains 10nM M13mp18 scaffold, 50nM staple mixture, 5mM tris acetate (pH 7.4), 5mM magnesium acetate and 0.5mM EDTA. The folding of the frame origami was carried out by heating the mixture to 95°C for 5 minutes followed by gradual temperature decrease to 20°C. The temperature was then further decreased down to 4°C until use.

#### DNA origami purification

All purified DNA origamis were stored in the DNA origami storage buffer composed of 10mM tris acetate, 10mM magnesium acetate, 1mM EDTA unless otherwise stated. All origamis were stored at 4°C and analysed within 2 weeks. All purification procedures were performed in 1.5ml centrifuge tubes unless otherwise specified.

##### SPRI beads selection

Two brands of commercially available Solid-phase reversible immobilization (SPRI) beads were tested throughout this study: HighPrep™ PCR Clean-up System (AC-60050; Magbio) and the SPRIselect (B23318; Beckman Coulter). The same procedure was used for both SPRI beads.

Prior to purification, the bead solution was set on the bench at room temperature for 15 minutes. The beads solution was then vortexed for 30 seconds until a homogenous colour was observed. The 40μl folded DNA origami after the folding reaction was mixed with 32μl of the SPRI beads to achieve the 0.8X volume ratio (for the effect of other ratio, refer to the main text for more information). The solution was mixed until a homogenous colour via flicking on the tube or pipette mixing, followed by 10 minutes incubation at room temperature on the bench. The tube was then placed on a magnetic separation rack (commercially available or homemade with 3D printer) for 5 minutes, the magnet pulled the SPRI beads from the solution to form a pellet at the corner of the tube. The clear supernatant was then isolated and discarded by pipette without disturbing the beads pellet. Then, two washes with 500µl of 80% (v/v) were performed and all excess ethanol from the tube was then carefully removed with a P10 pipette. The tube with the beads pellet was removed from the magnetic separation rack and sat on bench for 5 minutes to air dry. The dry pellet was then resuspended in 40µl of the DNA origami storage buffer through pipette mixing to elute the DNA origami from the SPRI beads, followed by 5 minutes of incubation time. The tube was placed back on the magnetic separation rack and the magnet caused the beads to pellet at the corner of the centrifuge tube, a pipette was used to aspirate the supernatant and transferred to a PCR tube. The purified DNA origami would need to undergo a thermal de-clumping procedure (refer to main text for further information about the process). The PCR tube was heated to 50°C using a thermal cycler for 2 mins, followed by a gradual decrease of the temperature to 20°C at the rate of 3°C/min. The temperature was then further decreased down to 4°C until use.

#### S-400 HR spin column filtration

The Microspin™ S-400 HR spin columns (27-5140-01; Cytiva) was used to purify the DNA origami. The spin column was set-up according to the manufacturer instructions. After the removal of the storage solution from the column by centrifugation at 750xg for 1 minute, the spin column was equilibrated with DNA origami storage buffer by adding 200µl to the column followed by centrifugation at 750xg for 1 minute. The equilibration step was repeated for 2 more times, all the eluates collected up were discarded. The column was then transferred and inserted into a clean centrifuge tube, then 40µl of the folded DNA origami was added to the column and spun down at 750xg for 1 minute. The eluate from the column was collected which contained the purified DNA origami. The volume of the purified DNA origami was checked with a pipette, if the volume was less than 40µl, additional DNA origami storage buffer was added to bring the volume to 40µl.

#### Molecular weight cut-off (MWCO) membrane filtration

Two different 100 kDa MWCO filtration units were tested, the Amicon Ultra-0.5 Centrifugal Filter Unit (R0PB46526; Sigma-Aldrich) (referred to as MWCO-1 in the main text) and the Vivaspin® 500 Centrifugal Concentrator (VS0142; Sartorius) (referred to as MWCO-2 in the main text). Same purification procedure was used for both centrifugal units.

The centrifugal unit was set-up as instructed by the manufacturer, then 400µl of the DNA storage buffer was first added to the centrifugal cartridge followed by 40µl of the folded DNA origami solution. The unit was then centrifuged at 15,000xg for 10 minutes, the filtrate was discarded, 400μl of the DNA storage buffer was added to the centrifugal cartridge, this process was repeated 2 more times. Afterwards, the purified DNA origami solution was collected with a pipette. The volume of the purified DNA origami was checked with a pipette, if the volume was less than 40μl, additional DNA origami storage buffer was added to bring the volume to 40μl.

#### PEG precipitation

For the purification of DNA origami through poly(ethylene) glycol (PEG) precipitation, the method described by Stahl *et al.* [4] was used without any modifications. The 15% PEG precipitation buffer was prepared ahead of purification, it contained 15% (w/v) PEG 8000 (89510; Sigma-Aldrich), 5mM Tris, 1mM EDTA and 505mM NaCl, the components were mixed and incubated at 85°C for 10 minutes and placed on tube roller for 2 hours.

The 40μl folded DNA origami was further diluted down to 200μl by adding 160μl DNA origami storage buffer, then 600μl of a buffer containing 5mM Tris-HCl (pH 8.0), 1mM EDTA, 20mM $MgCl_2$, and 5mM NaCl was added to the diluted folded DNA origami solution. Then 800μl of the 15% PEG precipitation buffer was added to the diluted folded DNA origami solution, followed by mixing by flicking the tube. The mixture was centrifuged at 10,000xg for 15 minutes, the supernatant was carefully removed without disturbing the pellet. The pellet was then resuspended in 800μl of buffer containing 5mM Tris-HCl (pH 8.0), 1mM EDTA, 20mM $MgCl_2$ and 5mM NaCl, followed by the addition of 800μl 15% PEG precipitation buffer, mixed and then centrifuge at 10,000xg for 15 minutes, this step was repeated 2 more times. After the last centrifugation step and the removal of the supernatant, the pellet was resuspended in 40μl of DNA origami storage buffer.

#### Phase separation

For the purification of the DNA origami via phase separation method, the method described by Masukawa *et al.* [5] was used without modifications. Two buffers were required for this method. The first buffer was Dextran PEG buffer containing 8.3% (w/v) PEG 6000 (81255; Sigma-Aldrich), 0.83% (w/v) Dextran 200,000 (31398; Sigma-Aldrich), 40mM tris acetate, 1mM EDTA (15558042; Thermo fisher), and 5mM magnesium acetate at pH 8.3. The second buffer was the PEG buffer containing 8.3% (w/v) PEG 6000, 40mM tris acetate, 1mM EDTA and 5mM magnesium acetate at pH 8.3.

After the folding reaction, 320μl of the Dextran PEG buffer was added to a 40μl solution containing the folded DNA origami. The mixture was vortexed for 30 seconds. The mixture was then centrifuged at 1700xg for 1 minute, 240μl of the supernatant was removed and discarded, 240μl of PEG buffer was added to the tube. The tube was mixed again via vortex shaking for 30 seconds and centrifuge at 1700xg for 1 minute, 200μl of the supernatant was removed and discarded, 200µl of PEG buffer was added to the tube. The tube was vortexed again for 30 seconds and centrifuge at 1700xg for 1 minute, 280µl of the supernatant was removed and discarded, 280µl of PEG buffer. The remaining 40µl solution was considered as the purified DNA origami.

##### Ethanol precipitation

For the purification of the DNA origami via ethanol precipitation, the method described by Lei *et al.* [6] was used without modifications. For this method, the precipitation buffer containing 60% (v/v) ethanol, 10mM Tris-HCl, 5mM $MgCl_2$ and 1mM EDTA at pH 8.0.

After the folding reaction, 40µl of the ethanol precipitation buffer was added to the folded DNA origami solution, the tube was then subjected to centrifuge at 4500xg for 30 minutes at room temperature. The supernatant was discarded and the pellet was resuspended in 40µl of the DNA origami storage buffer.

##### Size Exclusion Chromatography (SEC)

A custom packed Sephacryl S-500 HR column was used. The column was packed according to the manufacturer instructions. The column bed volume was 40 ml, and the column inner diameter was 16 mm, the resin Sephacryl S-500 HR (17061310; Cytiva) was packed inside the column at a compression ratio of 1.15X, the AKTA pure system was used to facilitate and monitor the process. The column was equilibrated with the DNA origami storage buffer by running the column through 2 column volumes of DNA origami storage buffer. 320µl of folded DNA origami solution was diluted to 1ml in DNA origami storage buffer, the diluted folded DNA origami solution was loaded onto the column through AKTA, the purification was carried out at flow rate XX, different fractions of eluate were collected based on the $A_{280}$ trace. Fraction corresponded to the DNA origami were pooled and concentrated first via 3,000 MWCO Vivaspin® 6 Centrifugal Concentrator (VS0691; Sartorius) and then via the 3,000 MWCO Vivaspin® 500 Centrifugal Concentrator (VS0191; Sartorius) down to 40 µl.

#### Automated purification via liquid handling robot

For the automated purification based on the usage of SPRI beads, the liquid handling system – Biomek $NX^P$ Automated Workstation (Beckman Coulter) was used. The DNA origami was folded inside a 96 well plate with each well containing 20µl of folded DNA origami mixture. The entire purification procedure including magnet beads separation, ethanol rinse, incubation and liquid transfer was set-up in the liquid handling system. Due to the volume limit of a 96 well plate, only 120µl of 80% ethanol was used to rinse the beads pellet. The final beads elution volume was 20µl to maintain the input and output volume consistency. The eluted plate from the liquid handling robot was then subjected to thermal de-clumping procedure as described above.

### DNA origami yield measurement

##### Absorption spectroscopy

The concentration of the purified DNA origami was measured via absorption spectroscopy using a NanoDrop™ 2000c Spectrophotometer (Thermo Fisher). The machine was blanked with the DNA origami storage buffer, 2µl of the sample was dotted onto the measurement platform, measurement was performed using an extinction coefficient of 33 mg/ml for $A_{260}$ = 1.

##### Fluorescence

The concentration of the DNA origami was also measured with a fluorescence-based method which allows the measurement of numerous samples in a 96 wells format. The QuantiFluor® ONE dsDNA System (E4871; Promega) was used to measure the concentration of the purified DNA origami after the automated purification step via pipette robot. The set-up and procedure were followed as instructed by the manufacturer, but with multiple changes to adapt for the DNA origami: 1. The 1X TE buffer was replaced with the DNA origami storage buffer; 2. Instead of the 48.5 kbp λ dsDNA provided by the kit to use as the calibration curve, a known mass (72, 36 and 18 ng) of the purified DNA origami (from S-400 HR spin column purification) was used.

The measurement was carried out in The Spark® multimode microplate reader (TECAN) with excitation at 360 nm and emission at 535 nm. The plate was shaked for 10 seconds with double orbital mode prior to reading, each well was read 16 times at different position, the fluorescent average was calculated from 16 readings.

### Size selection of the DNA ladder

The GeneRuler 1 kb Plus DNA Ladder (SM1331; Thermo Fisher) was used with the HighPrep™ PCR Clean-up System to demonstrate the size selective behaviour of the SPRI beads. Briefly, three ratios were selected (0.4X, 1.0X and 4.0X), a total of 1000 ng of the DNA ladder was dilute in 20µl volume. The procedure of the SPRI beads selection follows the same procedure as above in DNA origami purification, except the thermal declump step was skipped.

### Agarose Gel analysis

For the agarose gel analysis of the DNA origami, a 0.8% agarose gel was used. The high purity agarose (16500500; Thermo Fisher) was mixed with 0.5X TBE, 10mM $MgCl_2$ buffer in an Erlenmeyer flask, the agarose was melted via microwave at full power for 45 seconds, and immediately poured into the casting module of the agarose gel electrophoresis system (Mini-Sub Cell GT Systems; Bio-Rad). Either a fixed mass (25 ng) of the purified DNA origami or a fixed volume (4 µl) of the purified DNA origami was used. The purified DNA origami was mixed with 2µl of the loading dye (B7025S; NEB) and the volume was brought up to a total of 8µl with DNA origami storage buffer. The M13mp18 ssDNA scaffold was prepared in the same way with a fixed mass of 25 ng for gel analysis. The gel was submerged in the 0.5X TBE, 10mM $MgCl_2$ buffer inside a cold room (4°C), the samples were loaded with pipette, 70V was applied for 90 minutes. After the agarose gel has finished running, the gel was transferred to a foil covered container with 30ml of 1X TAE buffer inside, then 10µl of the Diamond nucleic acid dye (H1181; Promega) was used and stained the gel for at least 30 minutes.

For the gel analysis of the DNA ladder, a 0.8% agarose gel was used. The agarose was mixed with 1X TAE, melted with microwave at full power for 45 seconds, and casted with the gel casting module as described above. The samples were prepared similarly to DNA origami but with a different loading dye (B7024S; NEB). The gel was ran at 60V for 60 minutes, followed by gel staining with Diamond nucleic acid dye as described above.

The gel imaging was carried out with the InGenius LHR Gel Doc System (Syngene).

Further analysis of the gel such as densitometry analysis was done via ImageJ.

### Origami-protein mixture clean up

DNA origami purification from excess functional molecules: Folded origami structures were purified using SPRI beads as described earlier. The concentration of origami solution was checked with absorption spectroscopy. 24 nM of C-reactive proteins (CRP; 140-11R; Lee-Bio) was added to 12 nM of purified 4FST origami. The clean up of the mixture follows the procedure described above.

### SDS-PAGE analysis

A gradient Sodium dodecyl sulphate polyacrylamide gel electrophoresis (SDS-PAGE) gel was prepared by first preparing a 4% and 20% polyacrylamide gel by mixing 1M Tris-base, 0.3% SDS, pH adjusted with HCl to 8.45 buffer with 30% acrylamide/bis-acrylamide (29:1), 1% ammonium persulfate and 0.1% Tetramethylethylenediamine (TEMED).

For the preparation of the gradient gel, 1 volume portion of the 4% gel mixture was aspirated with a 10 ml serological pipette with an electronic pipette controller, then the same pipette was used to aspirate 1 volume portion of the 20% gel mixture. The serological pipette was withdrawn from the gel buffer tube, an air bubble was then generated by gently press the aspiration button from the controller, the air bubble was allowed to raise. After the air bubble disappeared, the gradient gel mixture was dispensed to the narrow gap in the sandwiched 1.0 mm glass plates. 1 mm size comb was inserted, and the gel was allowed to solidify. 15 µl of samples were mixed with 6 µl of 3X sample loading dye (187.5 mM Tris-HCl, 6% SDS, 30% glycerol, pH 6.8) so that the final mixture contains a 1X loading dye buffer and 12 ng of materials. 1X cathode (0.1M Tris base, 0.1M Tricine, 0.1% SDS, pH 8.3; J60992.K2; Alfa Aesar) and 1X anode buffer (0.2M Tris-HCl, pH 8.9) were used for running the gel. 20 µl of sample was loaded into the wells after the gel was solidified. The gel was then run at 200 V for 30 minutes. Staining of the gel was done by silver staining kit (24641; Thermo scientific) following the manufacturer's instruction. The silver stained all organic components including proteins and DNA. The stained gel was visualised and imaged using InGenius LHR Gel Doc System (Syngene).

### Atomic Force Microscopy (AFM)

The freshly cleaved mica discs were pre-treated with 5µl of 10mM $NiCl_2$, then immediately 5µl of the DNA origami samples were deposited onto the mica by pipetting directly into the $NiCl_2$ solution and left on bench for 5 mins, the samples were topped up with 100µl of DNA origami storage buffer. The DNA origami samples were imaged using a Bruker Dimension Fastscan (Santa Barbara, CA, USA) with Fastscan-D-ss or ScanAsyst-Fluid+ cantilevers containing a Si tip. The imaging was carried out via PeakForce tapping with ScanAsyst™ liquid imaging mode via Nanoscope software. All images acquired have a pixel resolution of 1024 X 1024, images were analysed with Nanoscope analysis 1.9.

### Intact origami percentage calculation

To calculate the percentage of the intact origamis from AFM images, the following rules were used.

- Structures having at least one side 80 nm considered as origami tile.
- Structures with at least half of one triangle or all combined missing is considered as a deformed tile.
- Only structures fully in AFM image frame is considered.
- Shape of a square as outline of structure only considered intact.
- The structure should have 2 parallel line adjacent to each other to be considered as intact.

Based on these rules, the table below summarises the number of origamis counted for each method, and the number of intact origamis.

**Supporting Table 1. The quantification of origami structures from AFM images for each method.**

| *Method Name* | *Intact origami* | *Damage origami* | *Total Origami Counted* |
|---|---|---|---|
| *0.8X SPRI* | 133 | 13 | 146 |
| *S-400 HR* | 54 | 8 | 62 |
| *MWCO-1* | 54 | 55 | 108 |
| *MWCO-2* | 24 | 1 | 25 |
| *PEG precipitation* | 38 | 1 | 39 |
| *Ethanol Precipitation* | 59 | 8 | 67 |
| *Phase Separation* | 21 | 2 | 23 |
| *SEC* | 79 | 14 | 93 |

### Streptavidin functionalised origami percentage calculation

To calculate the percentage of the streptavidin functionalised origami from AFM images, the following rules were used.

- Structures having at least one side 80 nm considered as origami tile.
- Only structures fully in AFM image frame is considered.
- Streptavidin on deformed origami is not considered.
- If two origami structures are linked with one streptavidin, it is counted as one.

Based on these rules, the table below summarises the number of origamis counted for the two methods

**Supporting Table 2. The quantification of streptavidin functionalised origami from AFM images for each method.**

| *Method Name* | *Streptavidin functionalised origami* | *Total Origami Counted* |
|---|---|---|
| *0.8X SPRI* | 102 | 120 |
| *S-400 HR* | 49 | 105 |

### Section 2: Supporting Figures

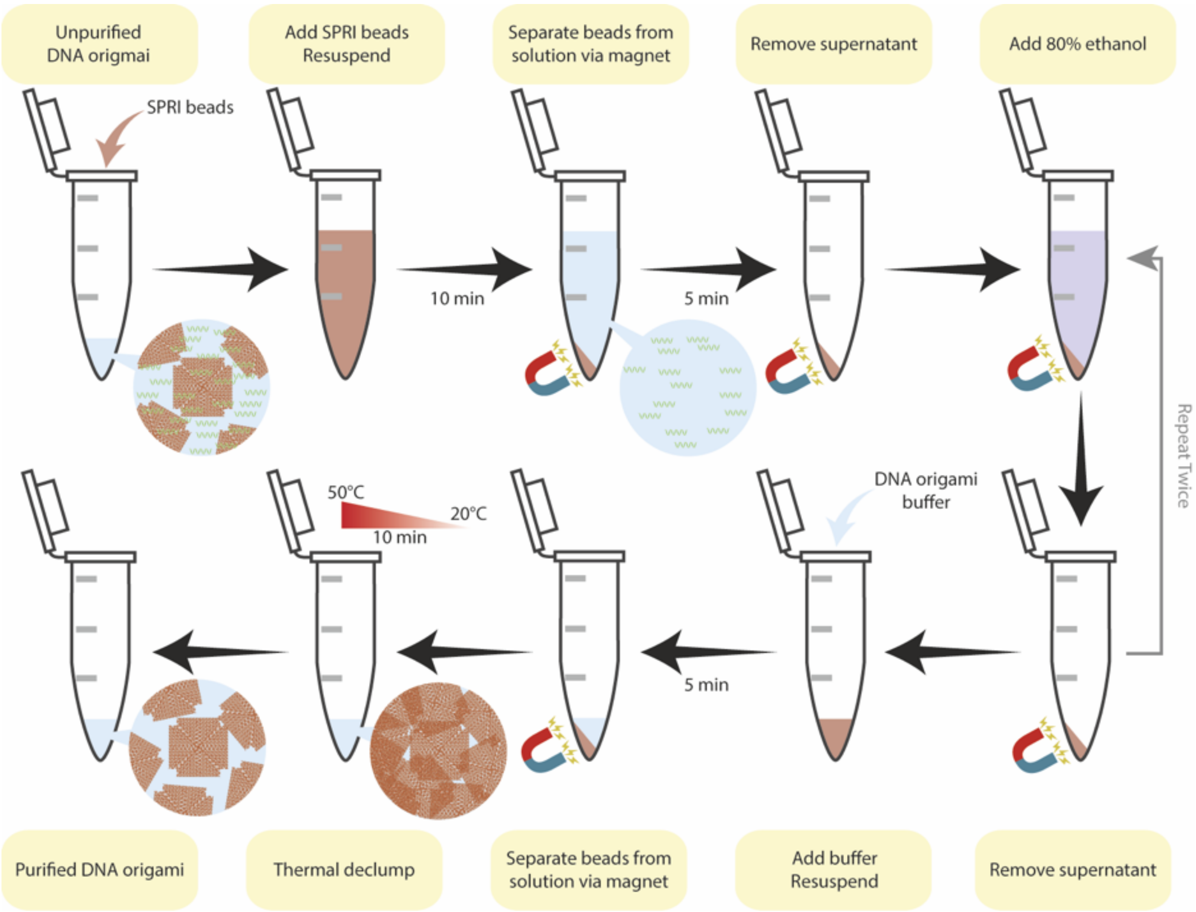

**Supporting Figure 1. The schematic illustration of the SPRI DNA origami purification routine.** The folded DNA origami after the folding reaction is then mixed with SPRI beads at a specific ratio, the SPRI beads with the DNA origami together form a beads pellet with the aid of a magnet, the excess staples in the supernatant are removed from the mixture. The beads pellet is washed with 80% ethanol twice to remove excess salt contamination from the SPRI beads buffer and the folding reaction buffer. The purified DNA origami can then be eluted in any buffer of choice (10 mM tris acetate, 10 mM magnesium acetate, 1 mM EDTA), the eluted DNA origami is subject to thermal declump procedure prior to finish.

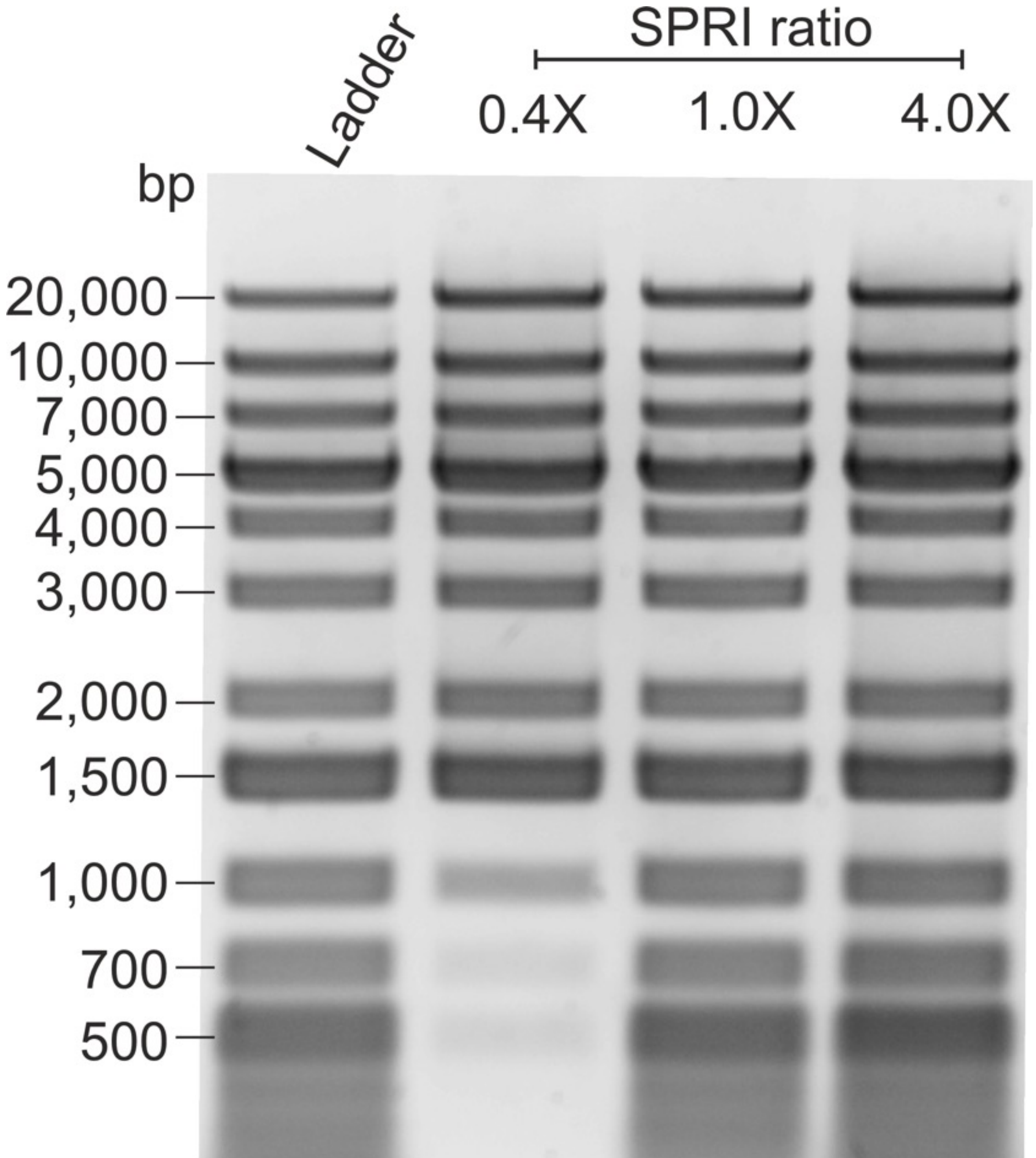

**Supporting Figure 2. The DNA size selection ability of the SPRI beads.** The DNA ladder used here was mixed with different ratio of SPRI beads. At low ratio of 0.4X, the beads selectively retain the higher molecular weight dsDNA from 1,000 bp onwards. At high ratio of 4.0X, the beads are not selective anymore and retain almost all the dsDNA regardless of the length. At 1.0X, only small dsDNA like 75 bp was excluded.

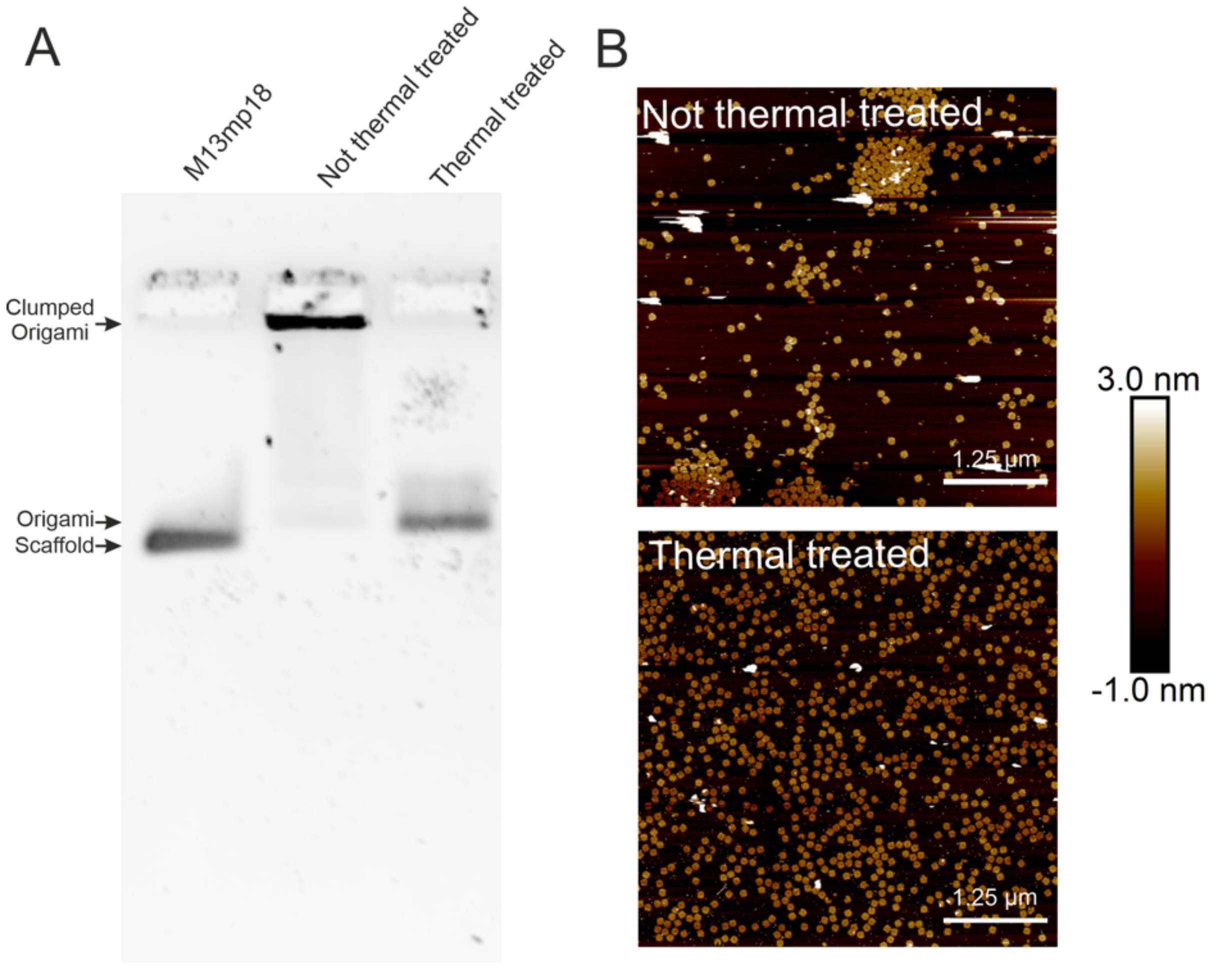

**Supporting Figure 3. The impacts of thermal de-clumping step in the SPRI beads based DNA origami purification method.** (A) The agarose gel shows that without the thermal de-clumping treatment step, the 4FST DNA origamis were not able to migrate through the agarose, suggesting aggregation of the materials. In contrast, the thermal de-clumping treatment of the same sample leads to the 4FST DNA origami to enter the agarose gel and migrate to the expected position. (B) AFM images of the thermal and not thermal treated 4FST DNA origami, as could be seen from AFM, the origami clumped together on the mica for the not thermally treated sample, the thermally treated origami dispersed evenly across the mica surface.

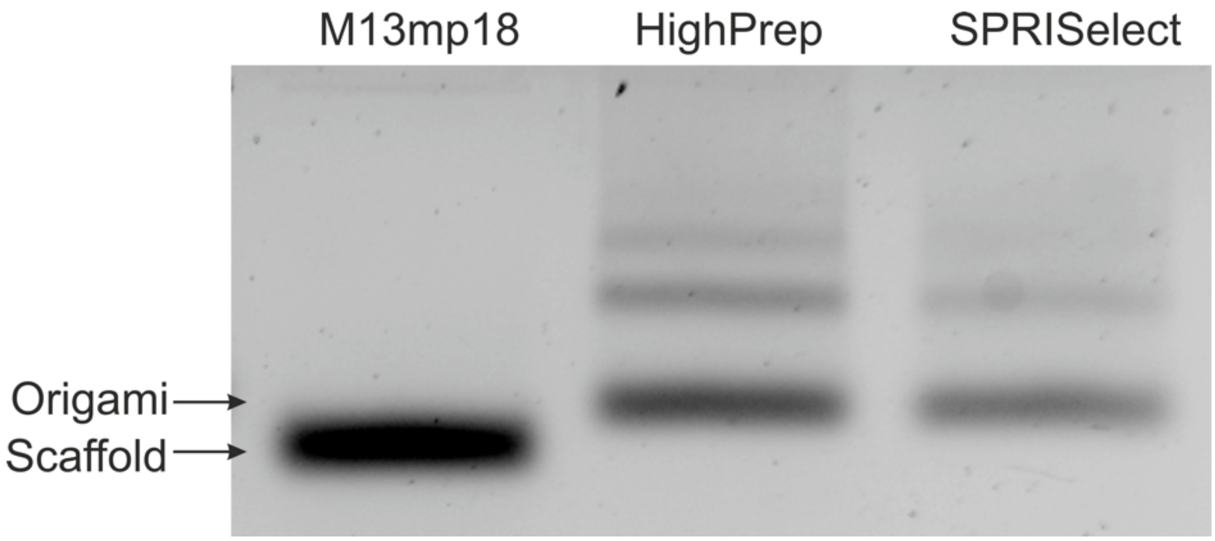

**Supporting Figure 4. Two different commercially available SPRI beads were tested.** Based on gel analysis, they have no differences in generation purification of the 4FST DNA origami. HighPrep SPRI beads would be used throughout the remaining study.

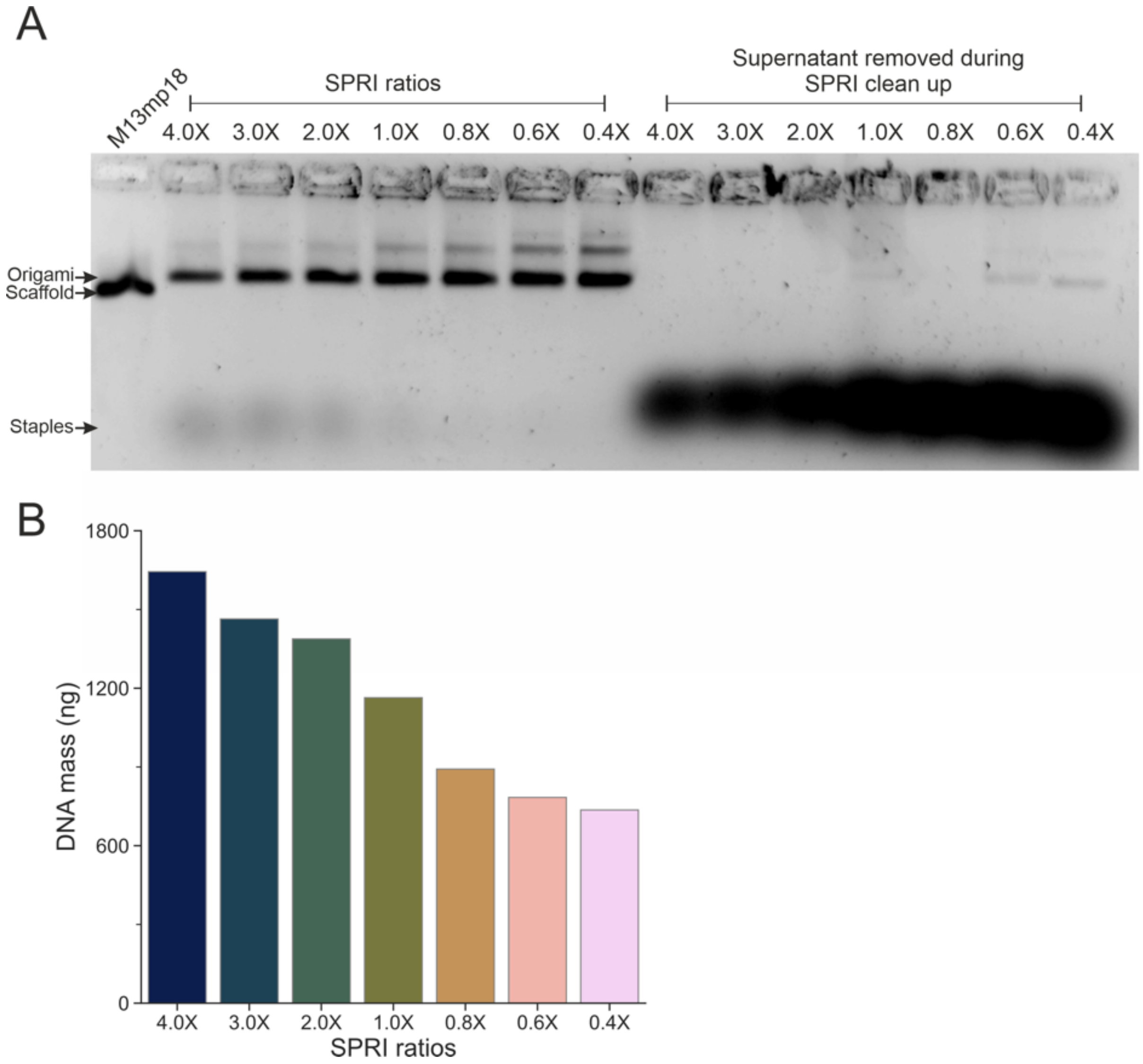

**Supporting Figure 5. The ratio screening of the SPRI beads on the purification of the 4FST DNA origami.** (A) The whole gel image from the main Figure 2. During the SPRI purification step, the supernatant was kept and analysed through agarose gel electrophoresis. As shown in the gel, the excess staples from the folding reaction could be found in the supernatant, suggesting that the SPRI beads efficiently retained the DNA origami and excluded the staples into the supernatant. For the analysis of the supernatant, a fixed volume of sample was added instead of fixed DNA mass. The stronger bands on the lower ratio indicate more staples, due to the mixing of the SPRI beads and folded DNA origami solution, the concentration of materials on the higher ratio size would be more diluted (*e.g.* 180 µl with 4.0X versus 56 µl with 0.4X), thus it was expected to observe more staples on the lower end due to a fixed volume of samples were loaded. (B) The yield of the DNA origami measured by absorbance at $A_{260}$. Although 4.0X had the highest yield, it could be seen from the gel that it was contaminated by the excess staples in the solution.

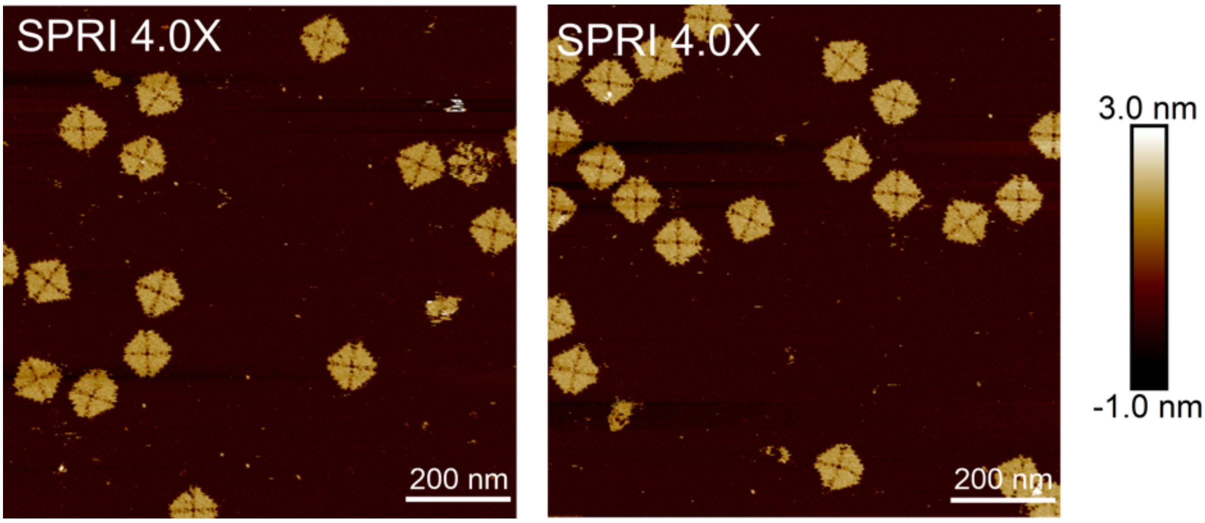

**Supporting Figure 6. The AFM images of the 4FST DNA origami purified with 4.0X SPRI beads ratio.**

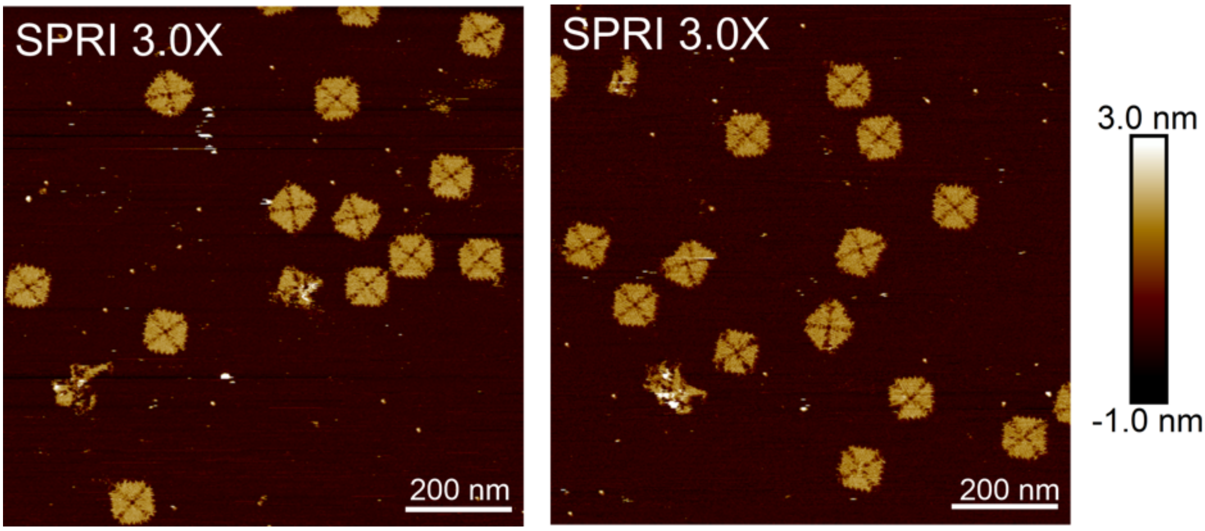

**Supporting Figure 7. The AFM images of the 4FST DNA origami purified with 3.0X SPRI beads ratio.**

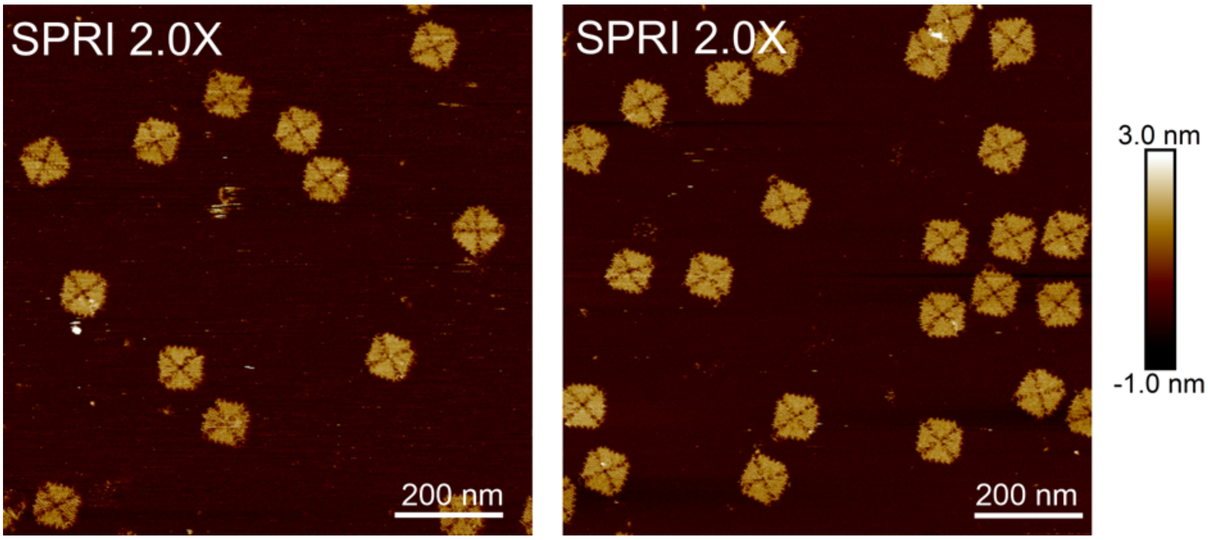

**Supporting Figure 8. The AFM images of the 4FST DNA origami purified with 2.0X SPRI beads ratio.**

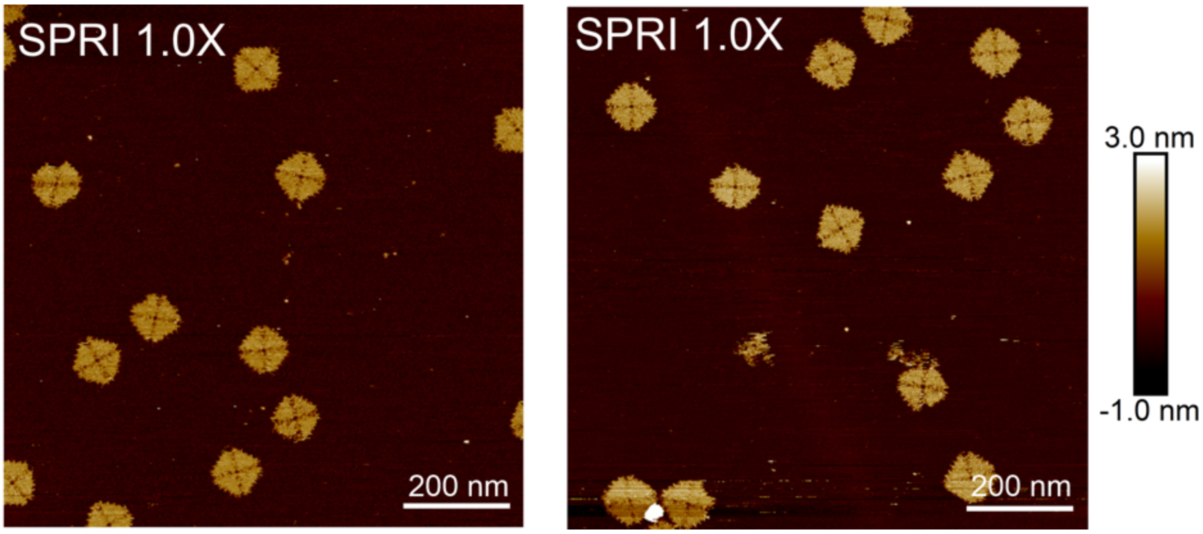

**Supporting Figure 9. The AFM images of the 4FST DNA origami purified with 1.0X SPRI beads ratio.**

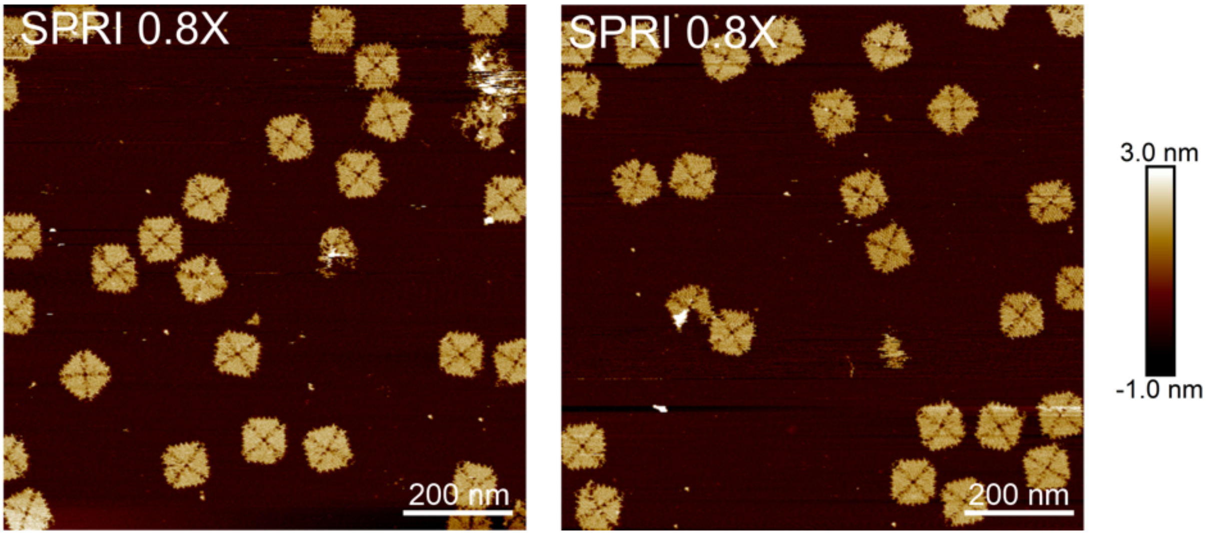

**Supporting Figure 10. The AFM images of the 4FST DNA origami purified with 0.8X SPRI beads ratio.**

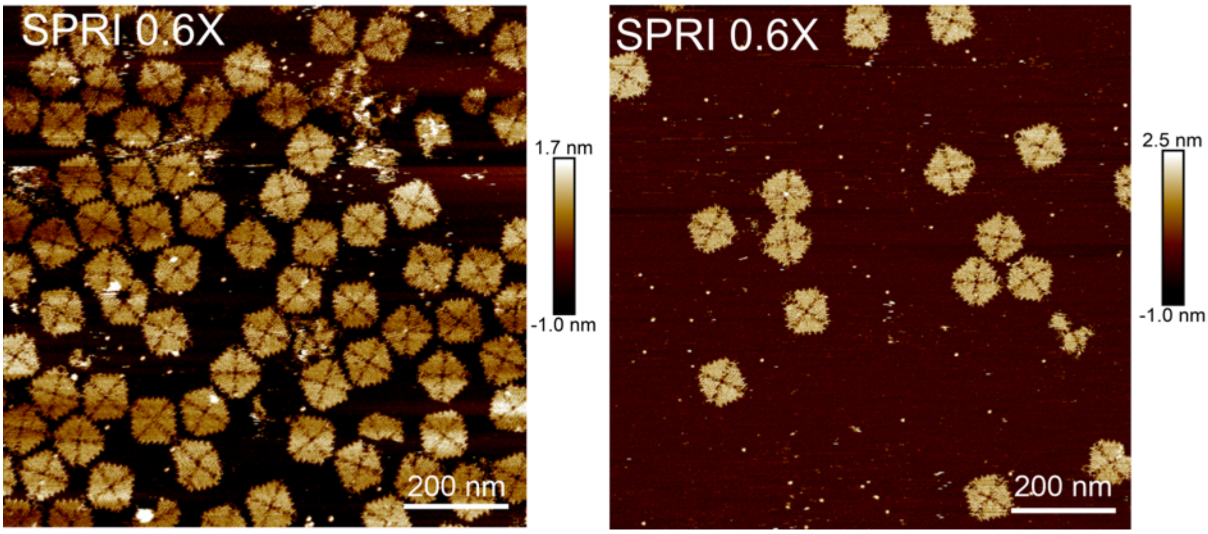

**Supporting Figure 11. The AFM images of the 4FST DNA origami purified with 0.6X SPRI beads ratio.**

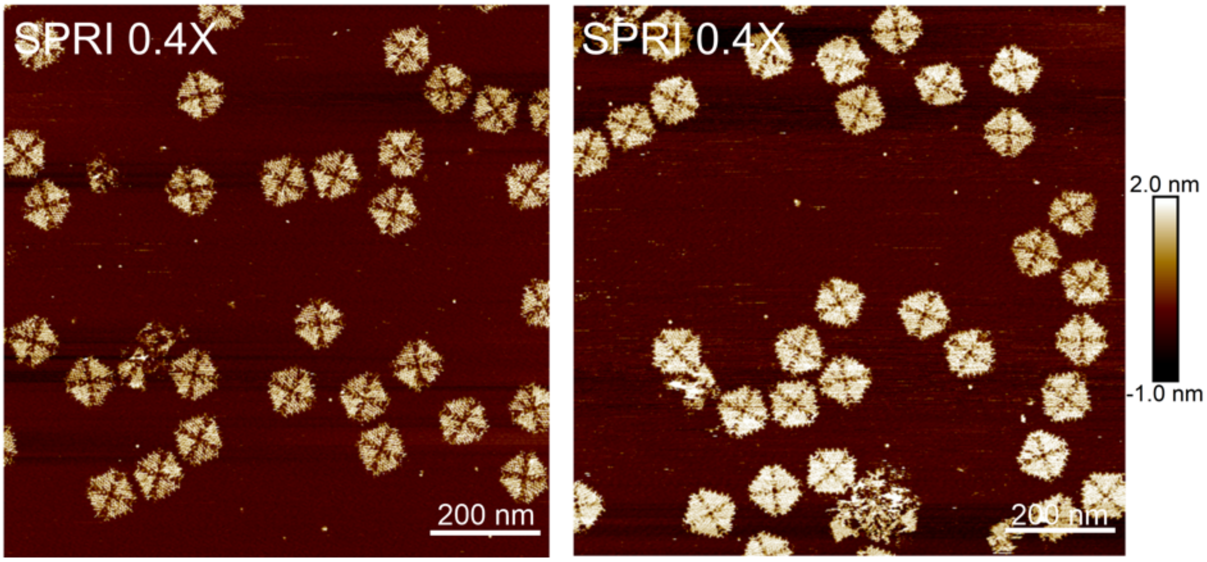

**Supporting Figure 12. The AFM images of the 4FST DNA origami purified with 0.4X SPRI beads ratio.**

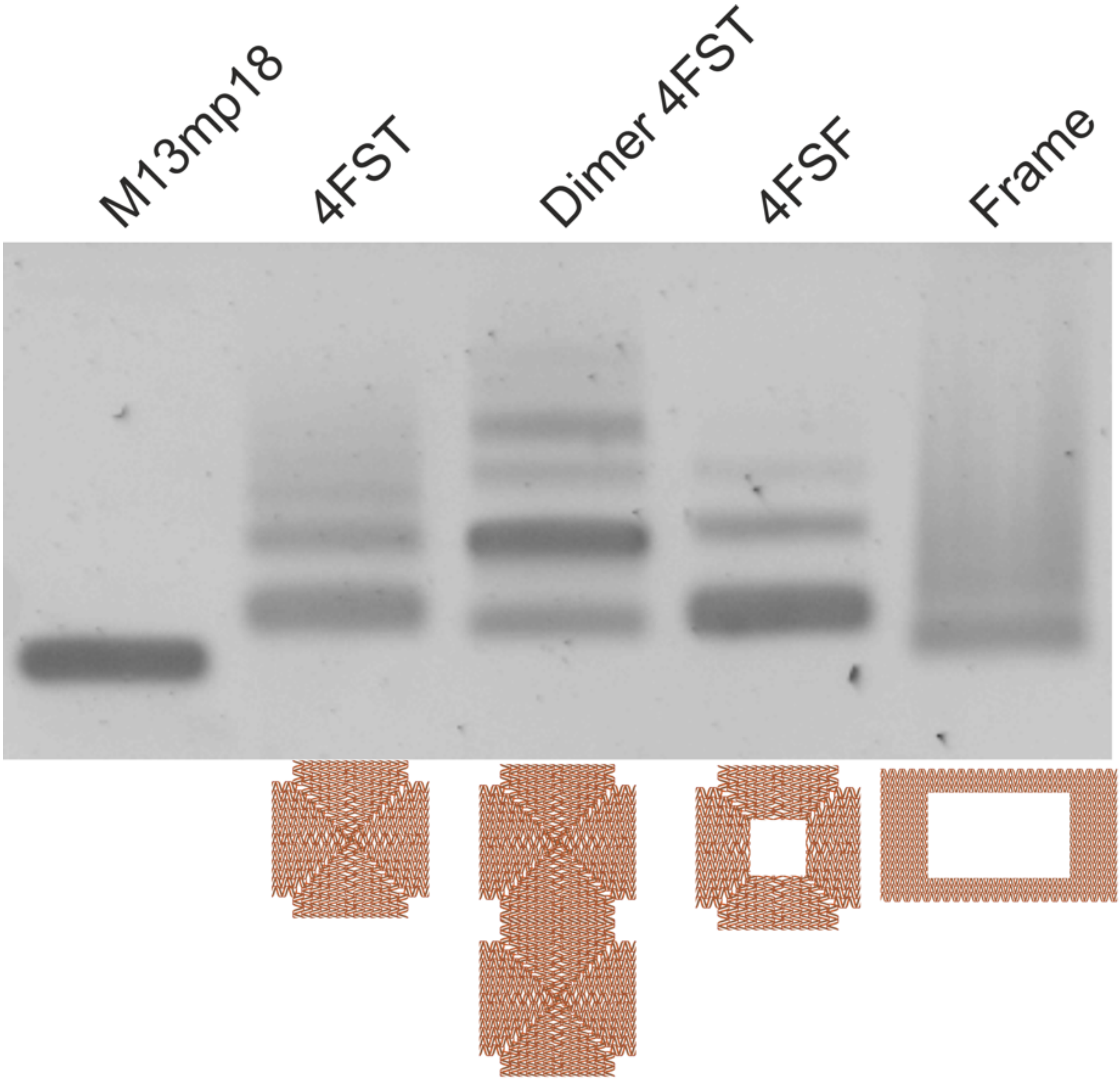

**Supporting Figure 13. Agarose gel electrophoresis analysis of the SPRI beads purified DNA origami structures.** Other DNA origami structures beside 4FST were purified with SPRI beads at the 0.8X ratio. The 4FST can dimerise to become the dimer 4FST. The 4FSF and frame origami both have a central cavity but of different dimensions. All structure migrates to the expected position on the agarose gel.

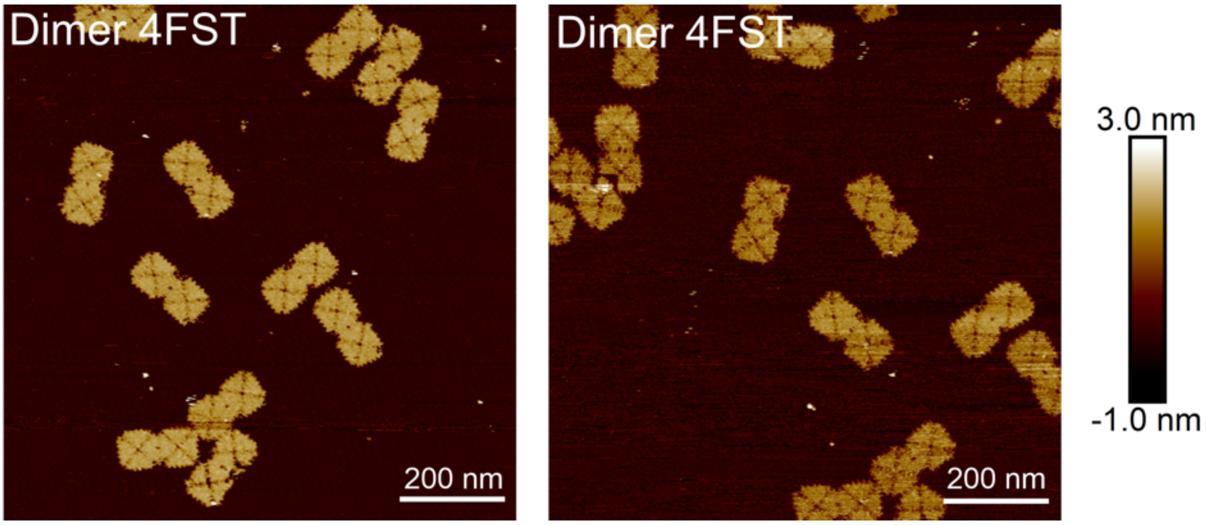

**Supporting Figure 14. The AFM images of the dimer 4FST DNA origami purified with 0.8X SPRI beads ratio.**

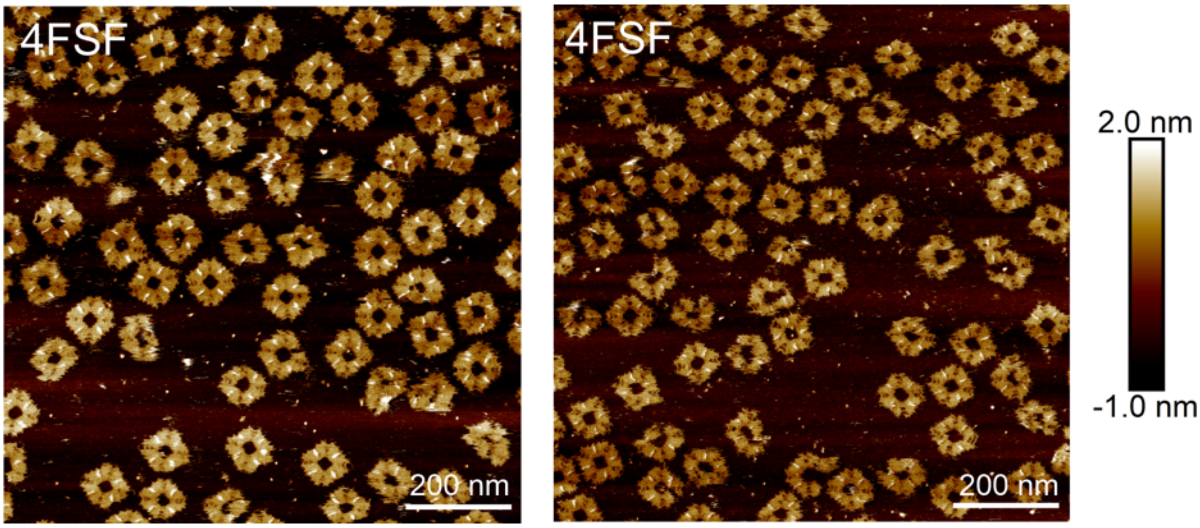

**Supporting Figure 15. The AFM images of the 4FSF DNA origami purified with 0.8X SPRI beads ratio.**

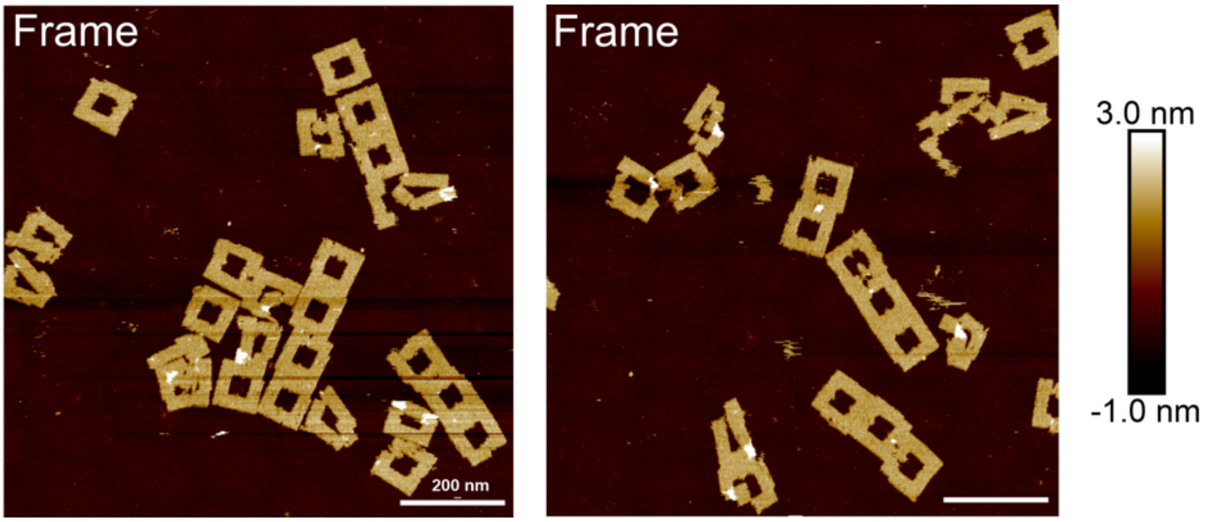

**Supporting Figure 16. The AFM images of the frame DNA origami purified with 0.8X SPRI beads ratio.**

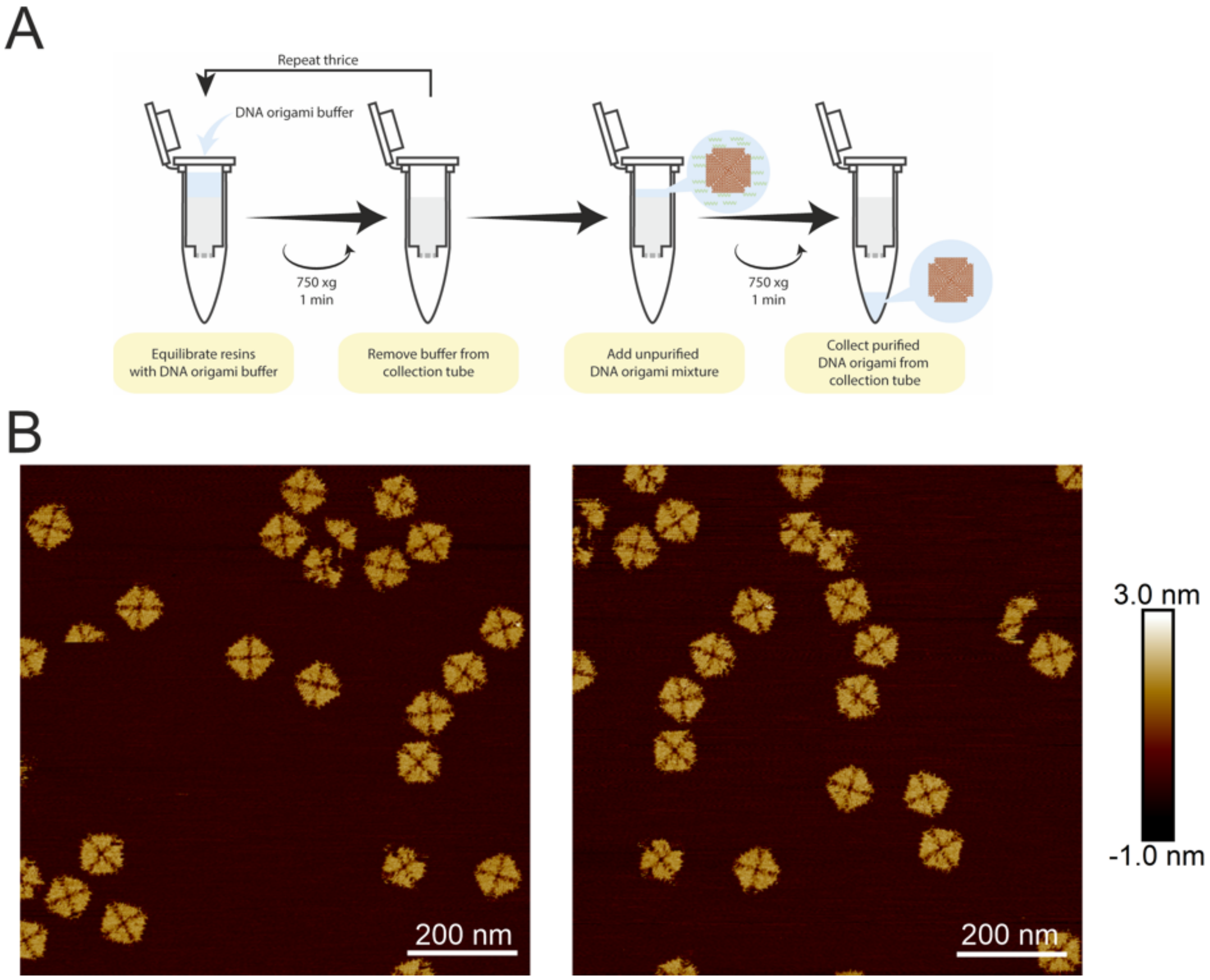

**Supporting Figure 17. The purification of the 4FST DNA origami through the S-400 HR spin column elution method.** (A) Schematic outline of the procedure. (B) The AFM images of the S-400 HR spin column elution purified 4FST DNA origami.

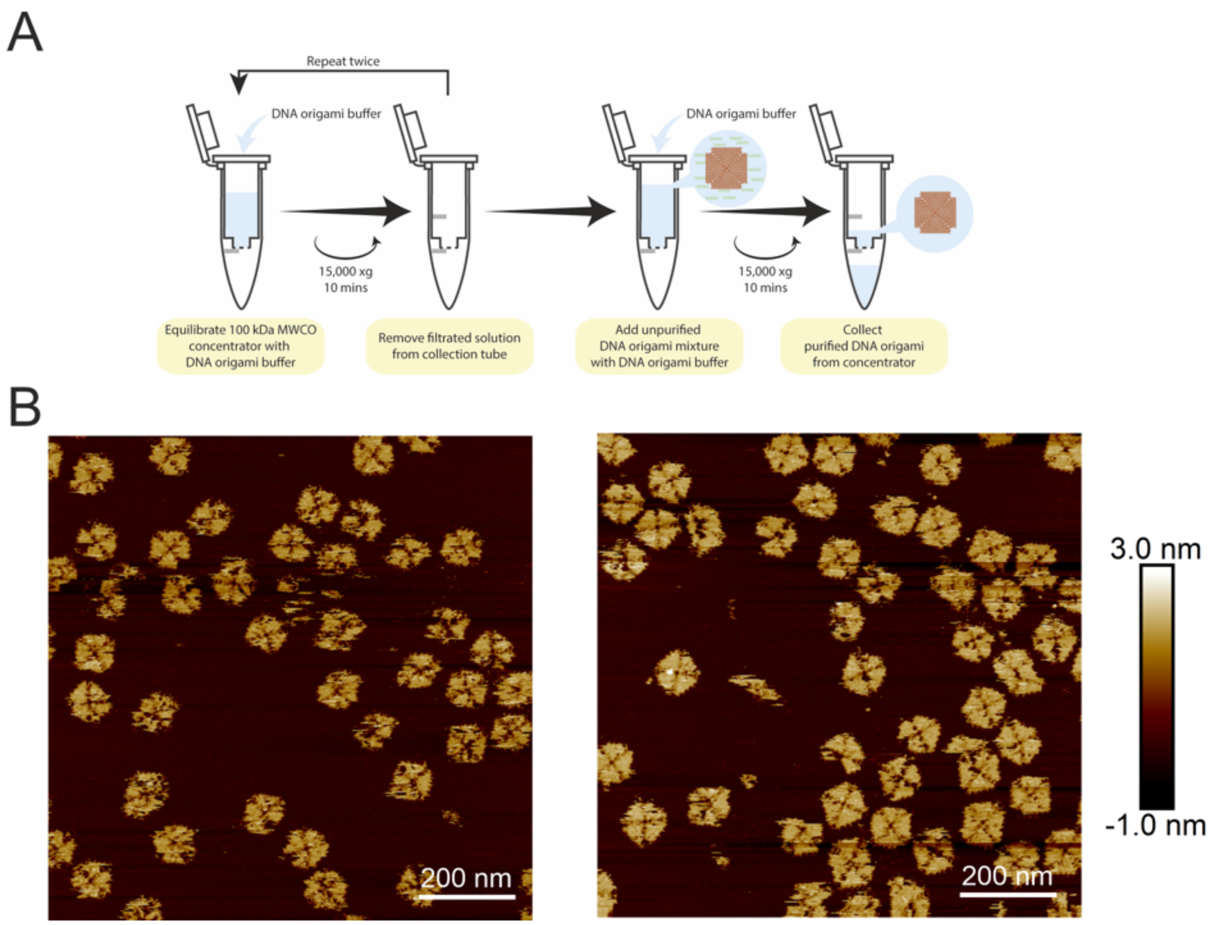

**Supporting Figure 18. The purification of the 4FST DNA origami through the 100 kDa MWCO membrane filtration (Amicon Ultra-0.5 Centrifugal Filter Unit, MWCO-1) method.** (A) Schematic outline of the procedure. (B) The AFM images of the 100 kDa MWCO purified 4FST DNA origami.

A

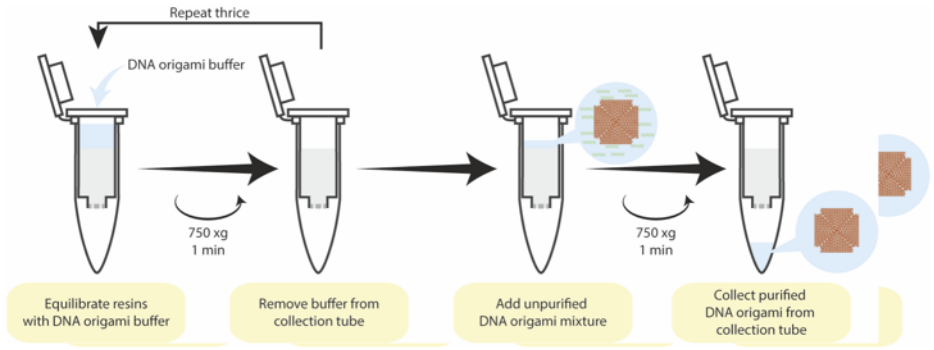

B

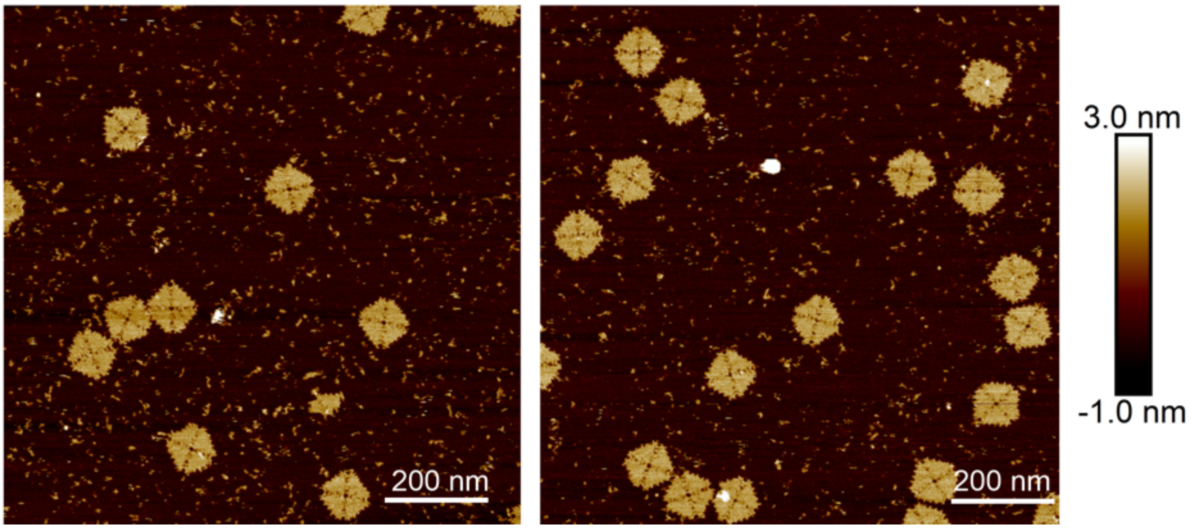

**Supporting Figure 19. The purification of the 4FST DNA origami through the 100 kDa MWCO membrane filtration (Vivaspin® 500 Centrifugal Concentrator, MWCO-2) method.** (A) Schematic outline of the procedure. (B) The AFM images of the 100 kDa MWCO purified 4FST DNA origami.

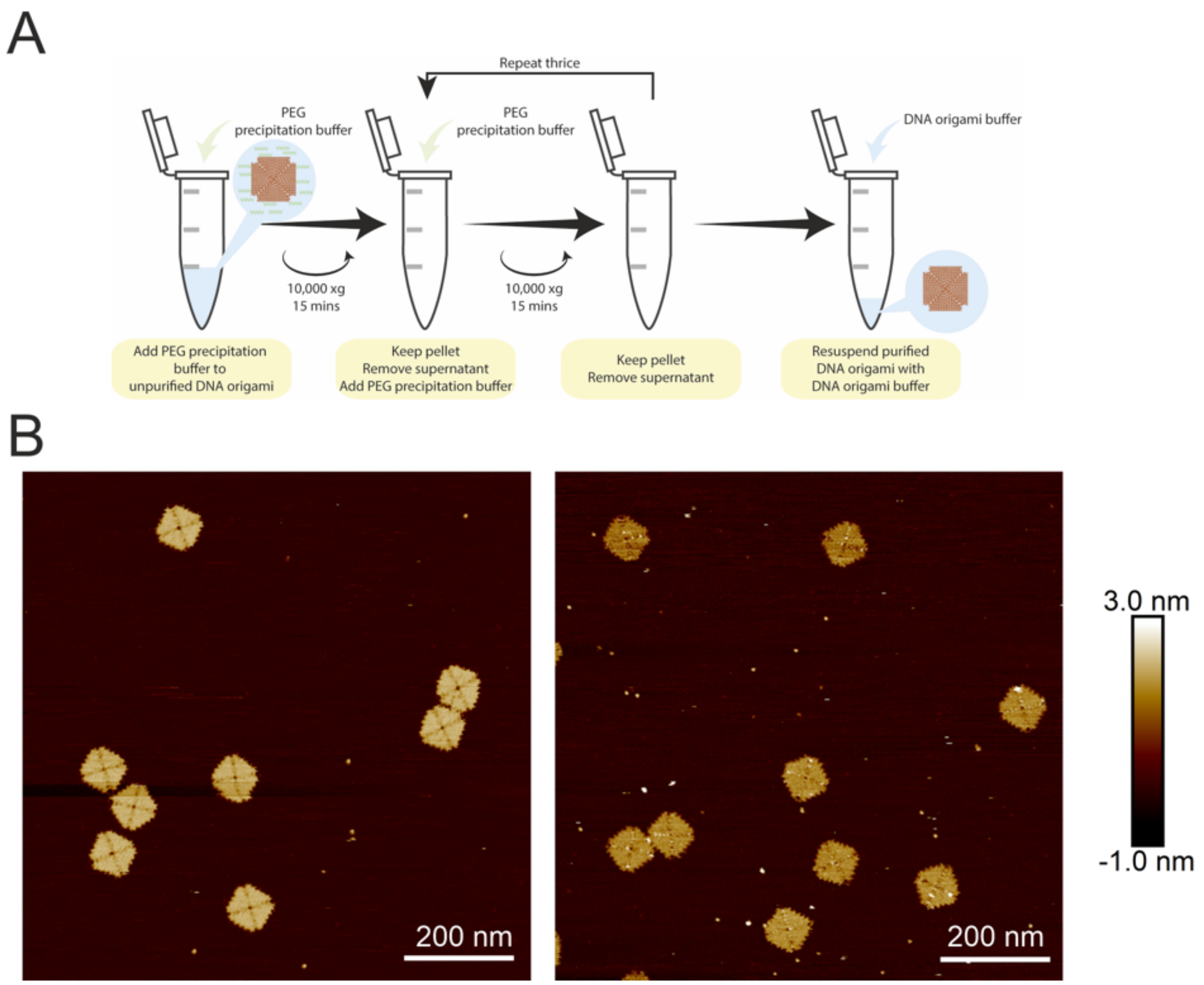

**Supporting Figure 20. The purification of the 4FST DNA origami through the PEG precipitation method.** (A) Schematic outline of the procedure. (B) The AFM images of the PEG precipitated 4FST DNA origami.

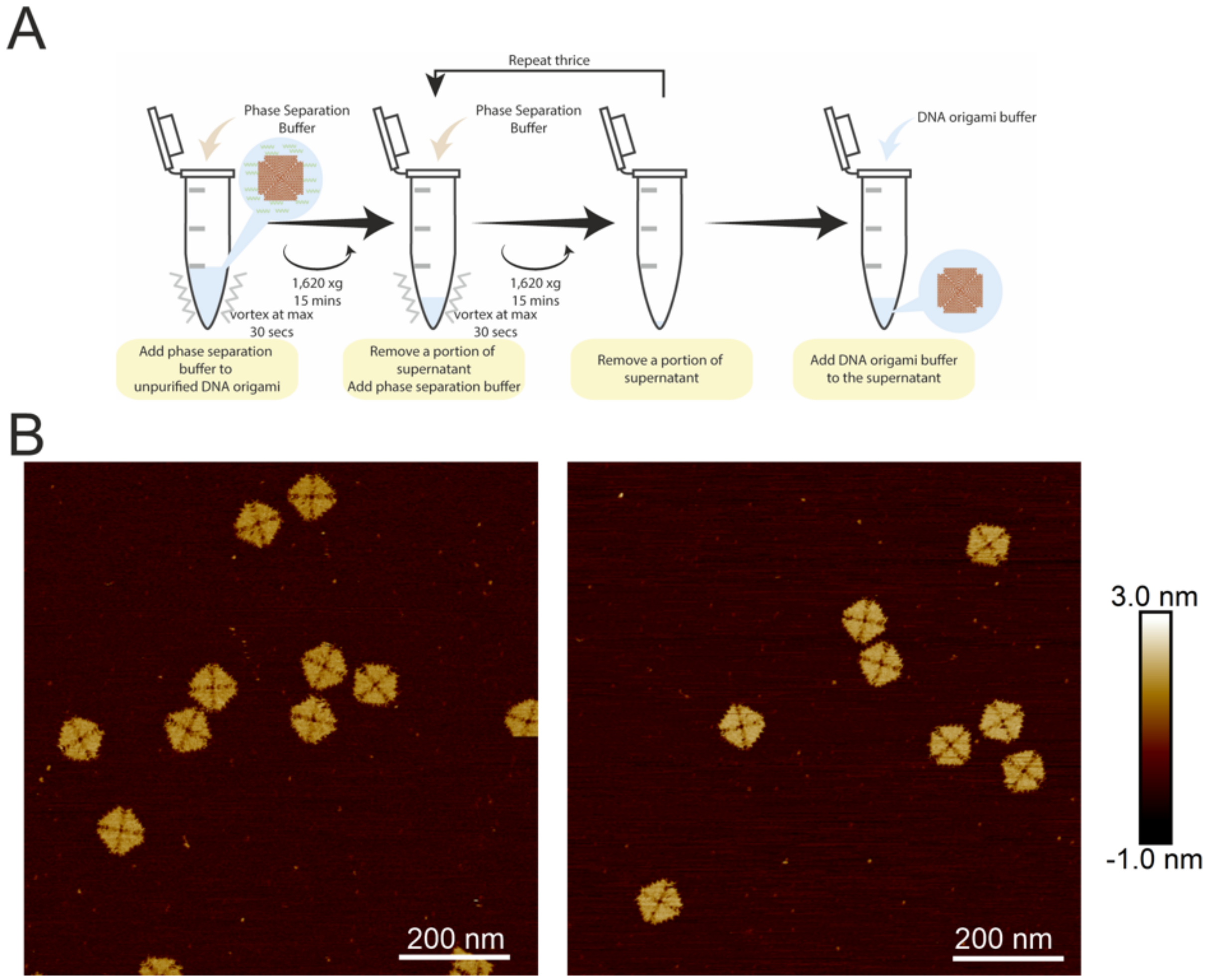

**Supporting Figure 21. The purification of the 4FST DNA origami through the phase separation method.** (A) Schematic outline of the procedure. (B) The AFM images of the phase separated 4FST DNA origami.

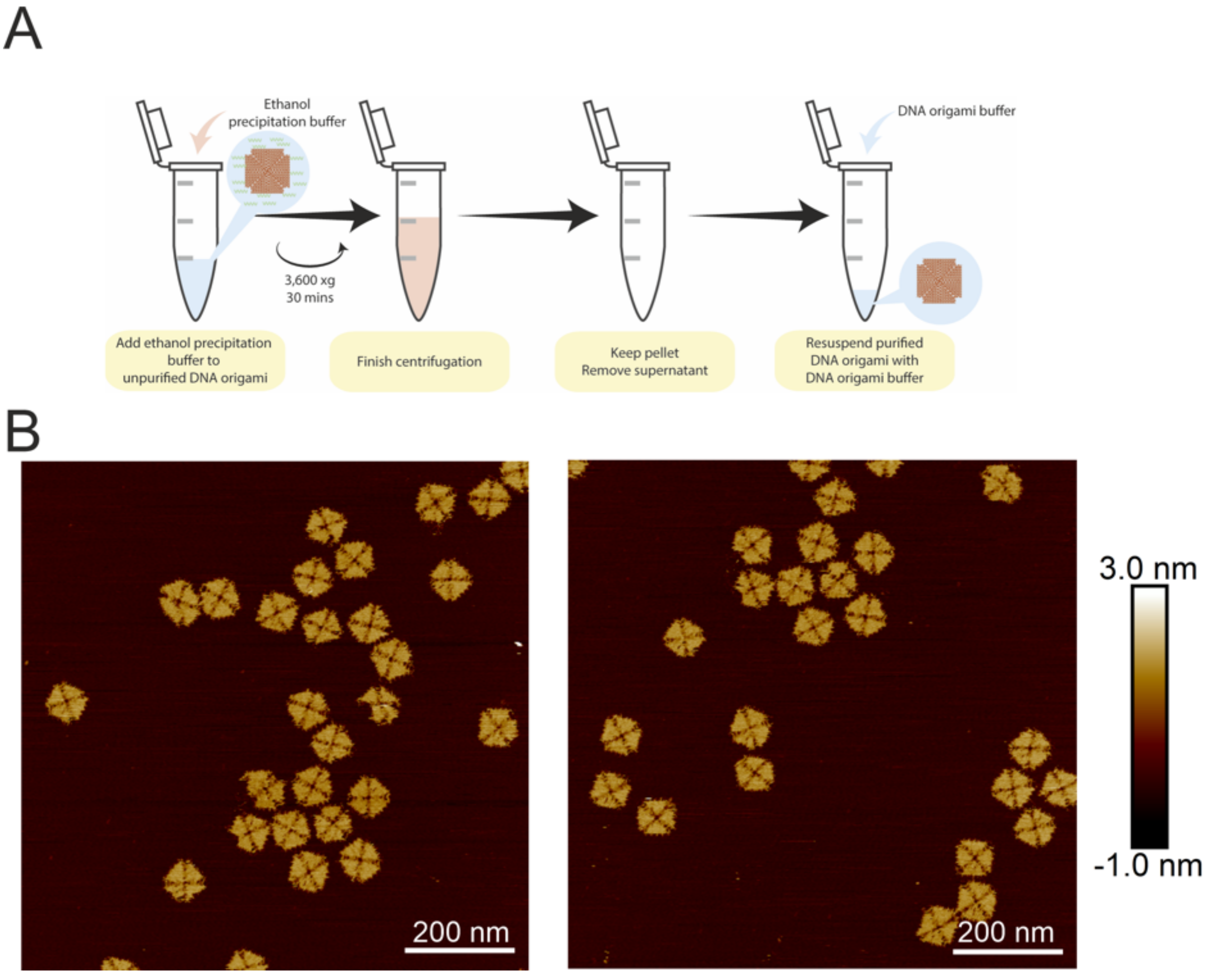

**Supporting Figure 22. The purification of the 4FST DNA origami through the ethanol precipitation method.** (A) Schematic outline of the procedure. (B) The AFM images of the ethanol precipitated 4FST DNA origami.

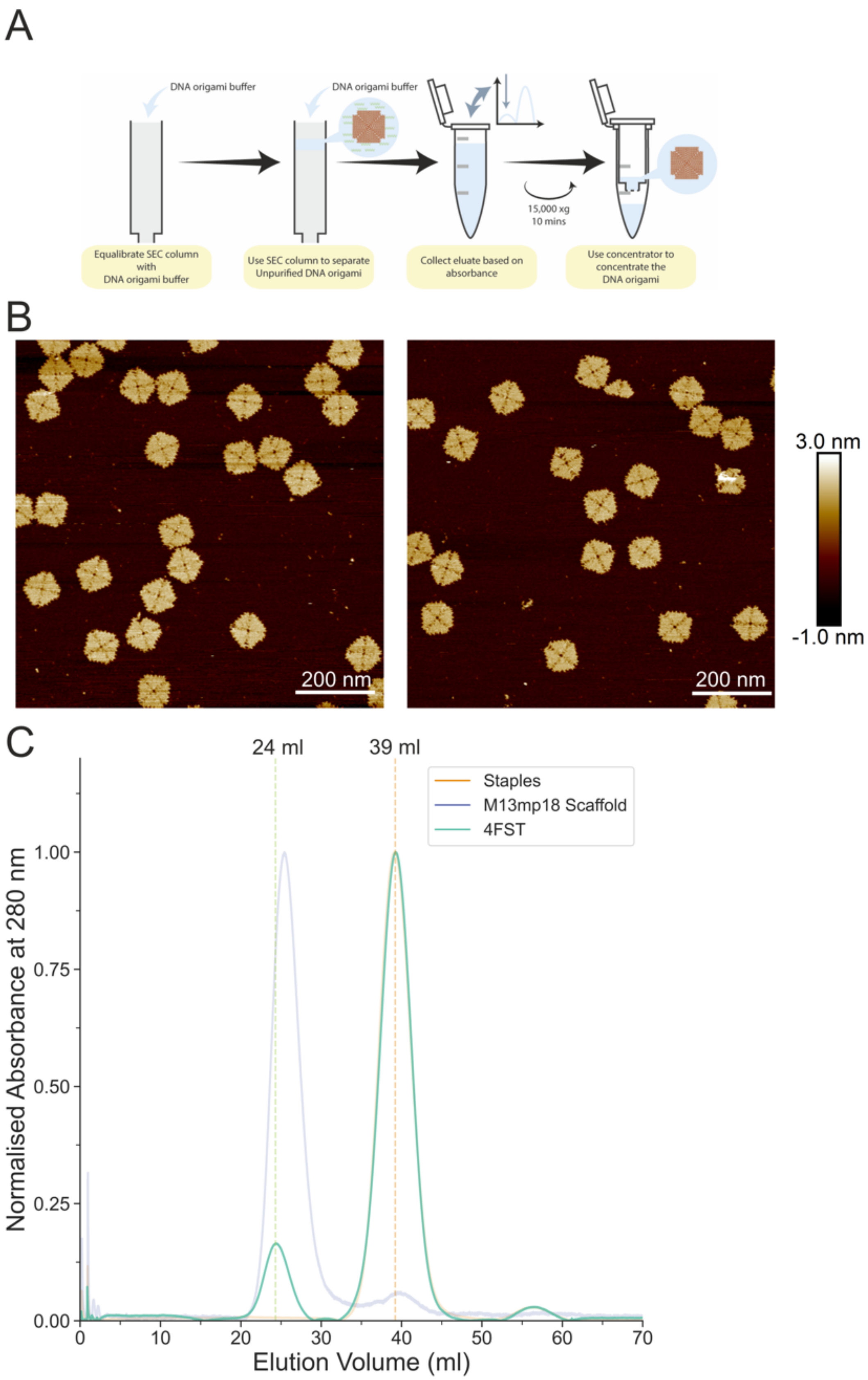

**Supporting Figure 23. The purification of the 4FST DNA origami through the SEC method.** (A) Schematic outline of the procedure. (B) The AFM images of the ethanol precipitated 4FST DNA origami. (C) The overlapped elution profile of staples, M13mp18 scaffold and the 4FST tile. The folded 4FST can be identified at 24 ml, the staples elute at 39 ml.

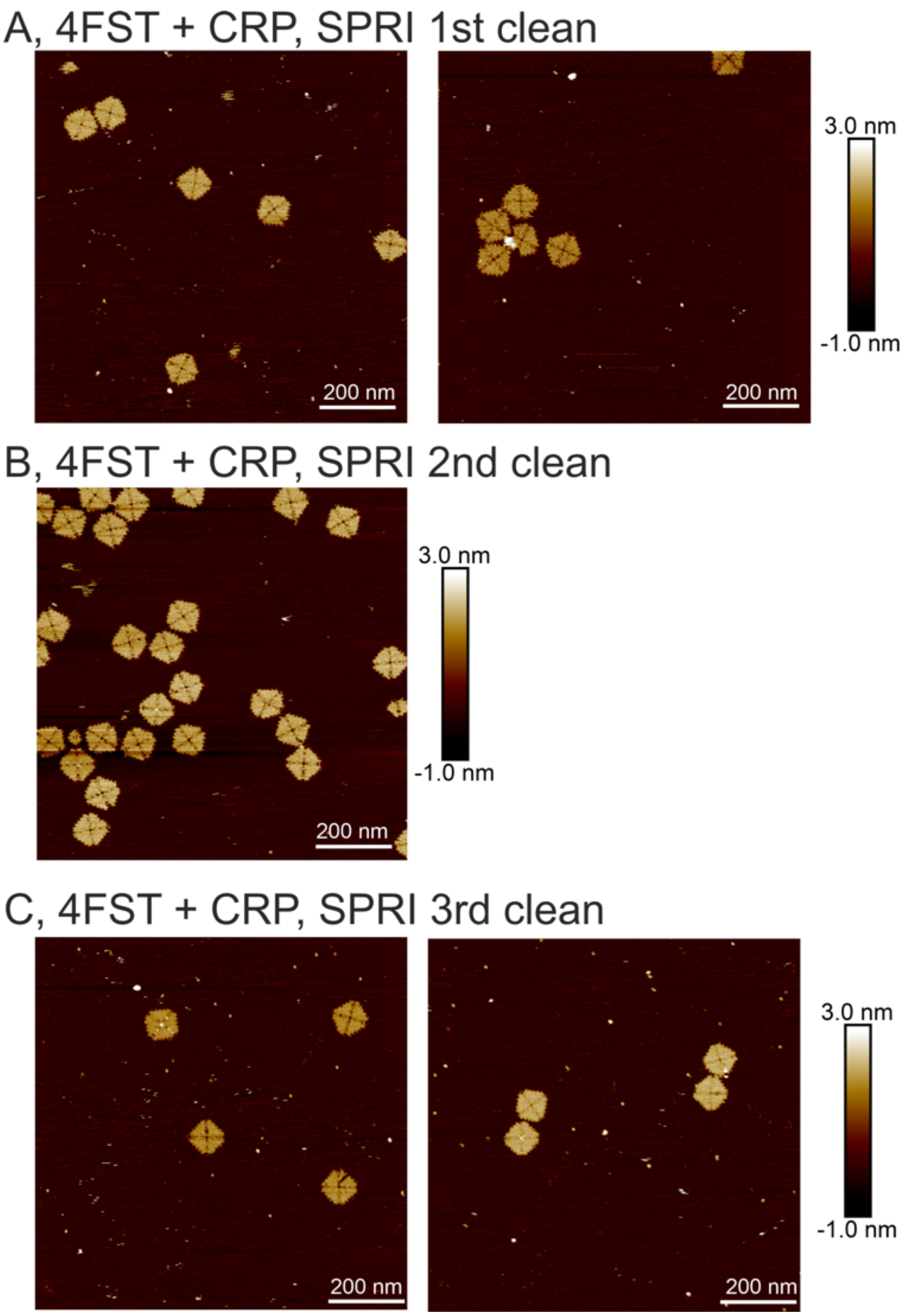

**Supporting Figure 24. AFM images of the SPRI beads purification of the 4FST to remove background CRPs.** (A) The 1$^{st}$ round of purification. (B) The 2$^{nd}$ round of purification. (C) The 3$^{rd}$ round of purification.

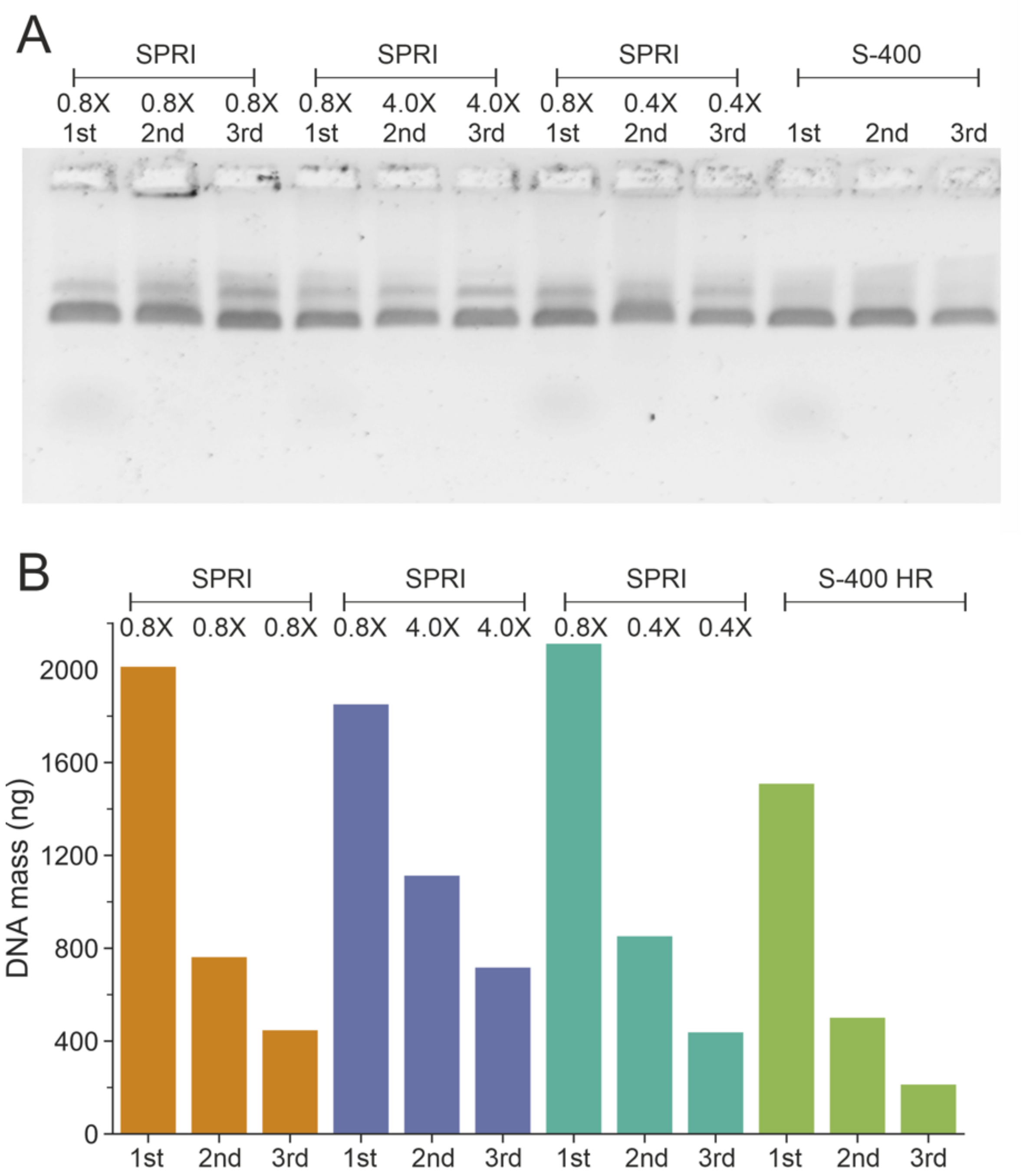

**Supporting Figure 25. Multiple rounds of purification comparison.** The 4FST DNA origamis were cleaned multiple rounds. (A) Gel analysis of the multiple rounds cleaned 4FST DNA origamis, all structures are intact and migrated to the expected distance. For the SPRI beads, a different ratio of beads were tested for the 2nd and 3rd round of purification to investigate whether this affect the structure and the final yield of the DNA origami. (B) The total DNA mass yield after multiple rounds of clean up measured by absorbance at $A_{260}$. Note, for this experiment a total of 80 µl origami reaction was used instead of 40 µl in other experiments. Thus the DNA mass is higher than previously shown.

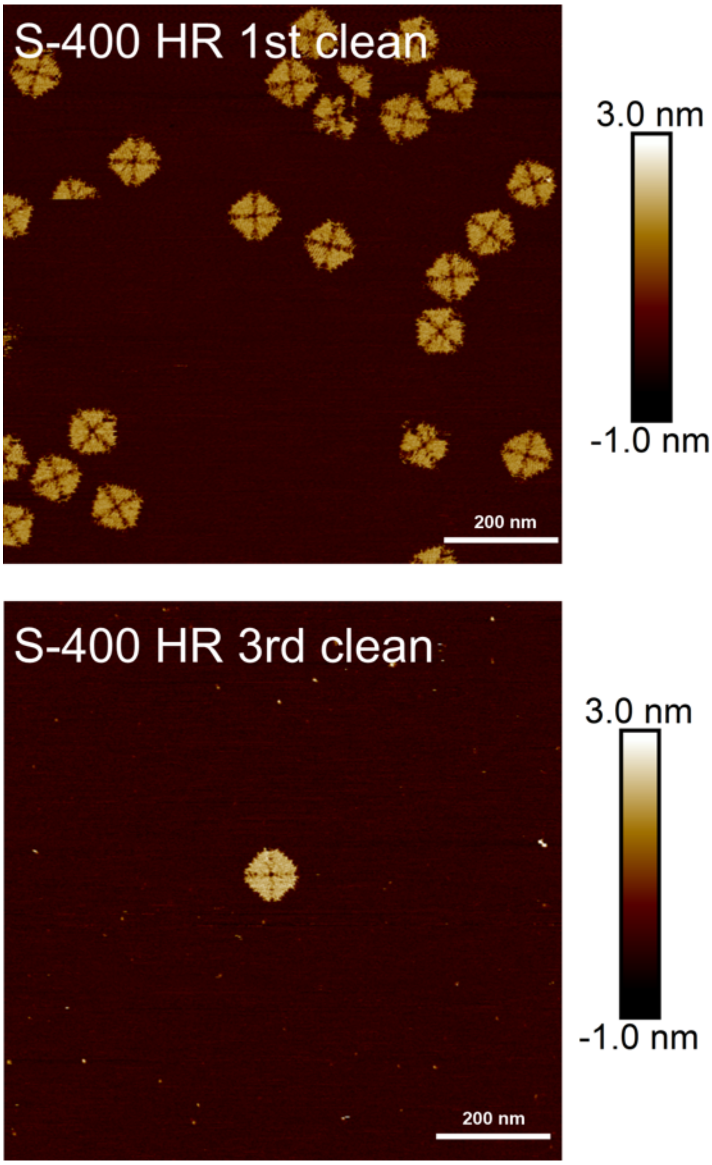

**Supporting Figure 26. The AFM images of the 4FST DNA origami after 3 rounds of S-400 HR spin column elution purification.** After 3 rounds of S-400 HR cleaned up, the concentration of the DNA origami dropped significantly, and origamis were hard to find through AFM.

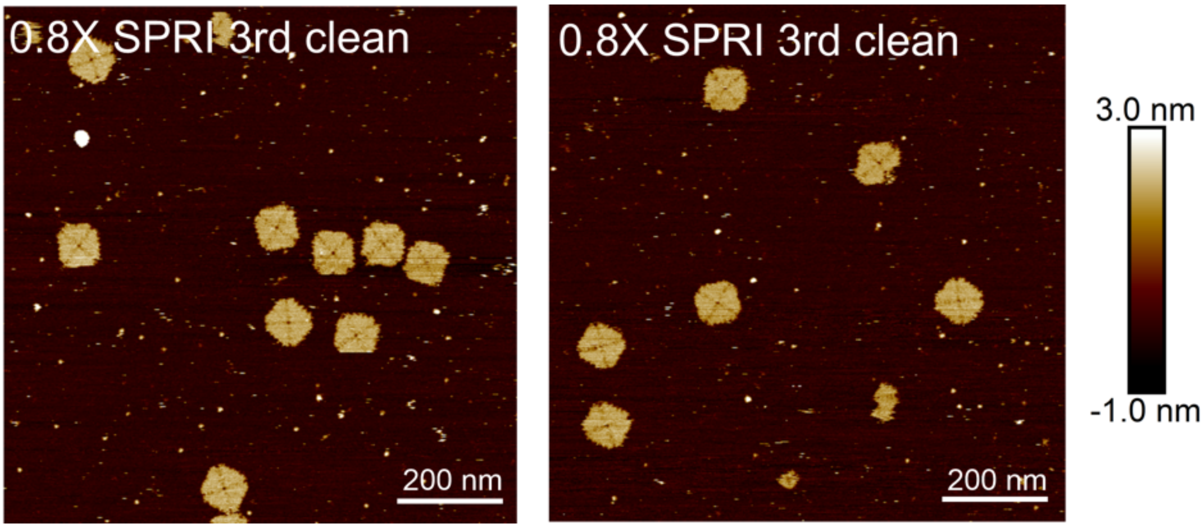

**Supporting Figure 27. The AFM images of the 4FST DNA origami after 3 rounds of SPRI beads purification.**

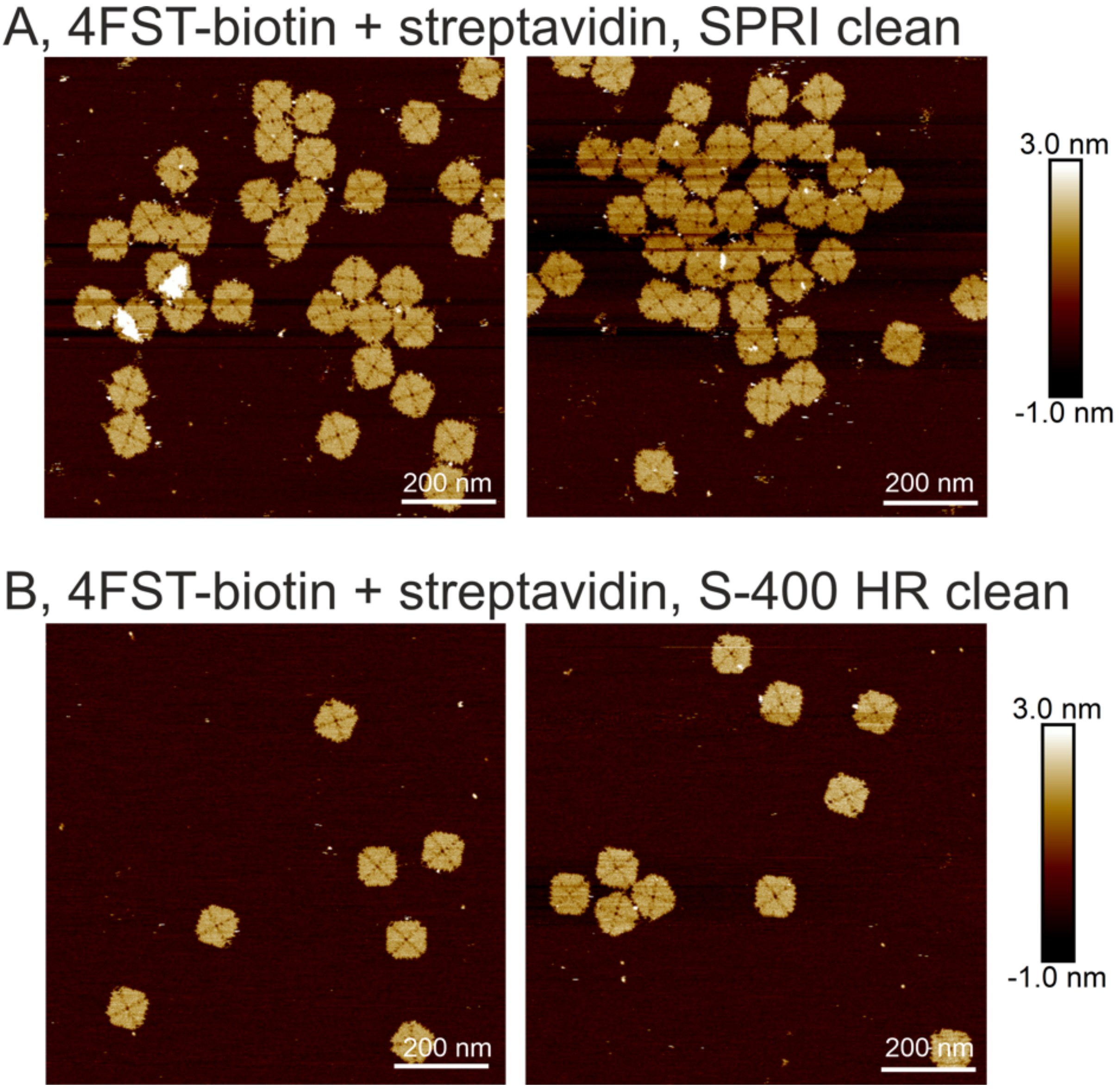

**Supporting Figure 28. Comparison between the SPRI and S-400 HR spin columns on the streptavidin functionalised biotinylated 4FST DNA origami.**

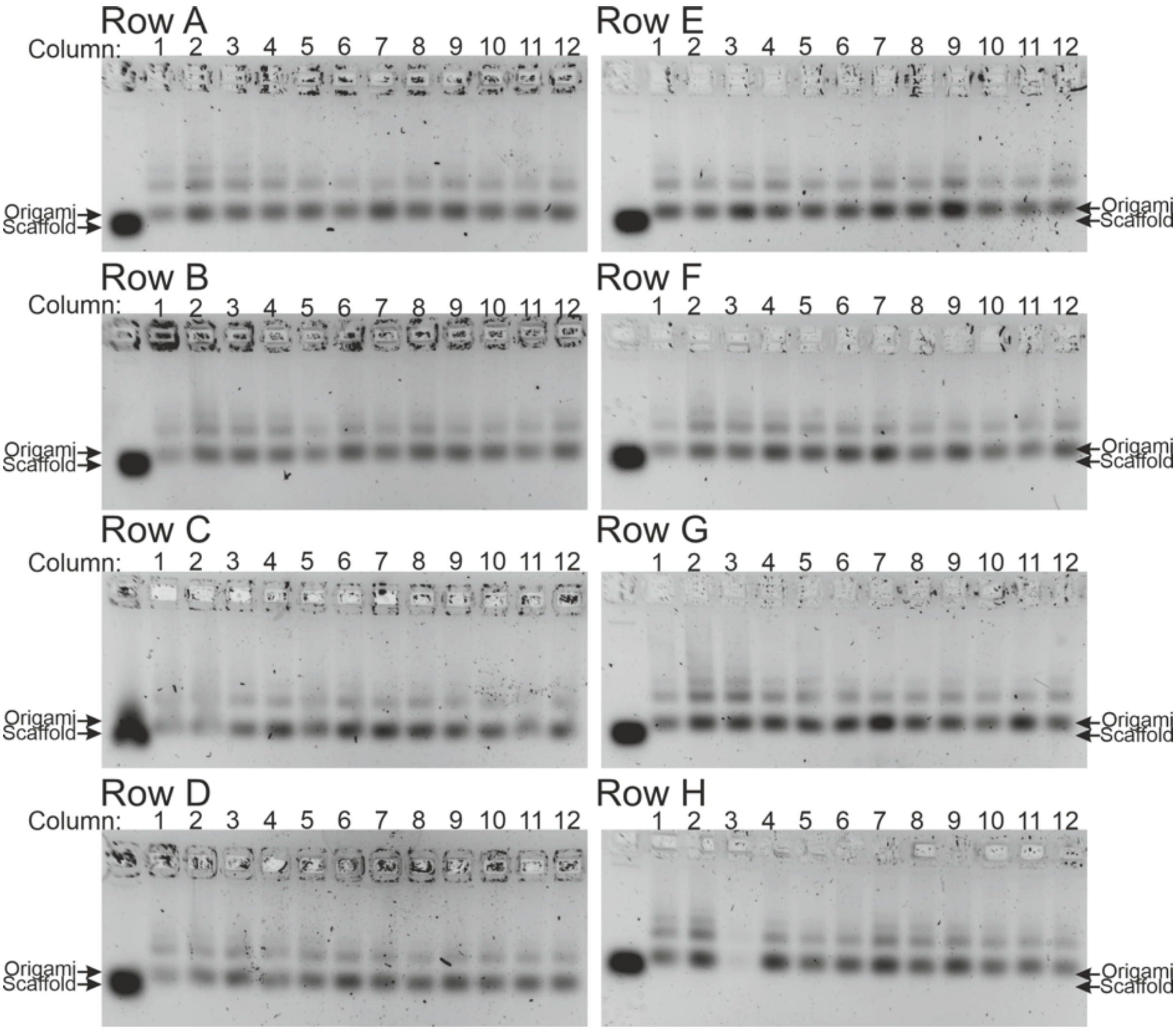

**Supporting Figure 29. Agarose gel electrophoresis of all the samples in the 96 well plate of Prep 1 after the SPRI based automated DNA origami purification via liquid handling robot.** The 4FST DNA origami can be identified in near 99% of the wells (95/96). This prep was performed with the HighPrep™ PCR Clean-up System

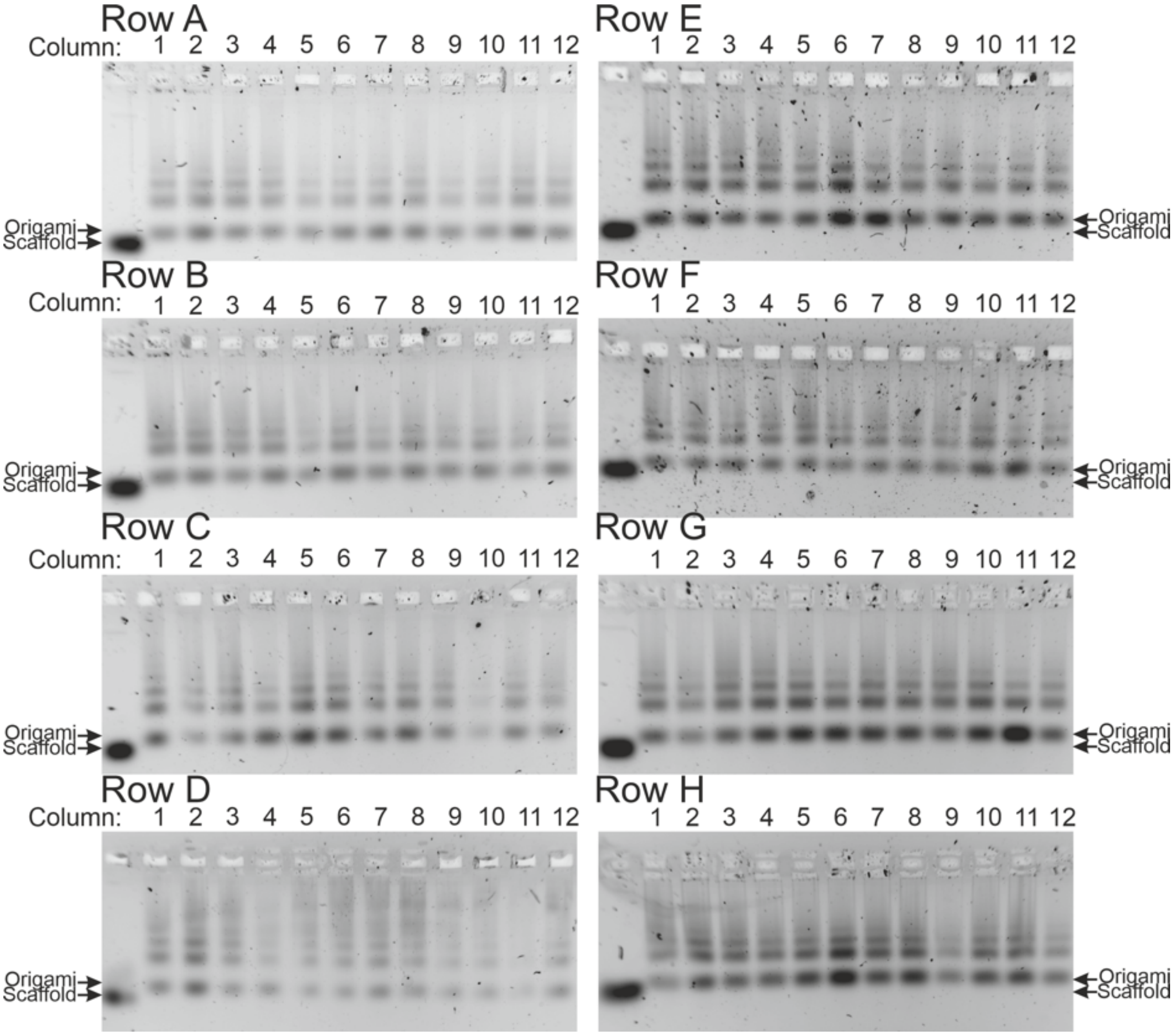

**Supporting Figure 30. Agarose gel electrophoresis of all the samples in the 96 well plate of Prep 2 after the SPRI based automated DNA origami purification via liquid handling robot.** The 4FST DNA origami can be identified in near 99% of the wells (95/96). This prep was performed with the SPRIselect.

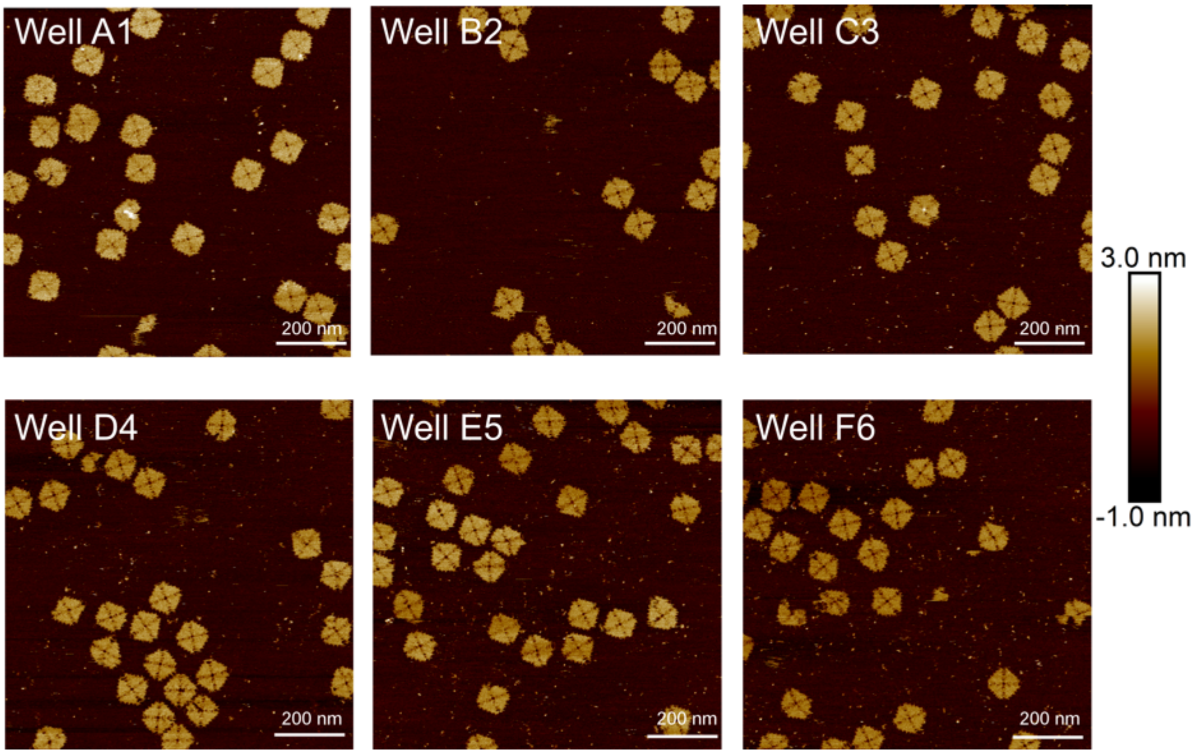

**Supporting Figure 31. AFM images of the SPRI based automated DNA origami purification via liquid handling robot.** 6 wells were selected (A1, B2, C3, D4, E5, F6) from the PCR plate of Prep 1 after the SPRI purification via the liquid handling robot.

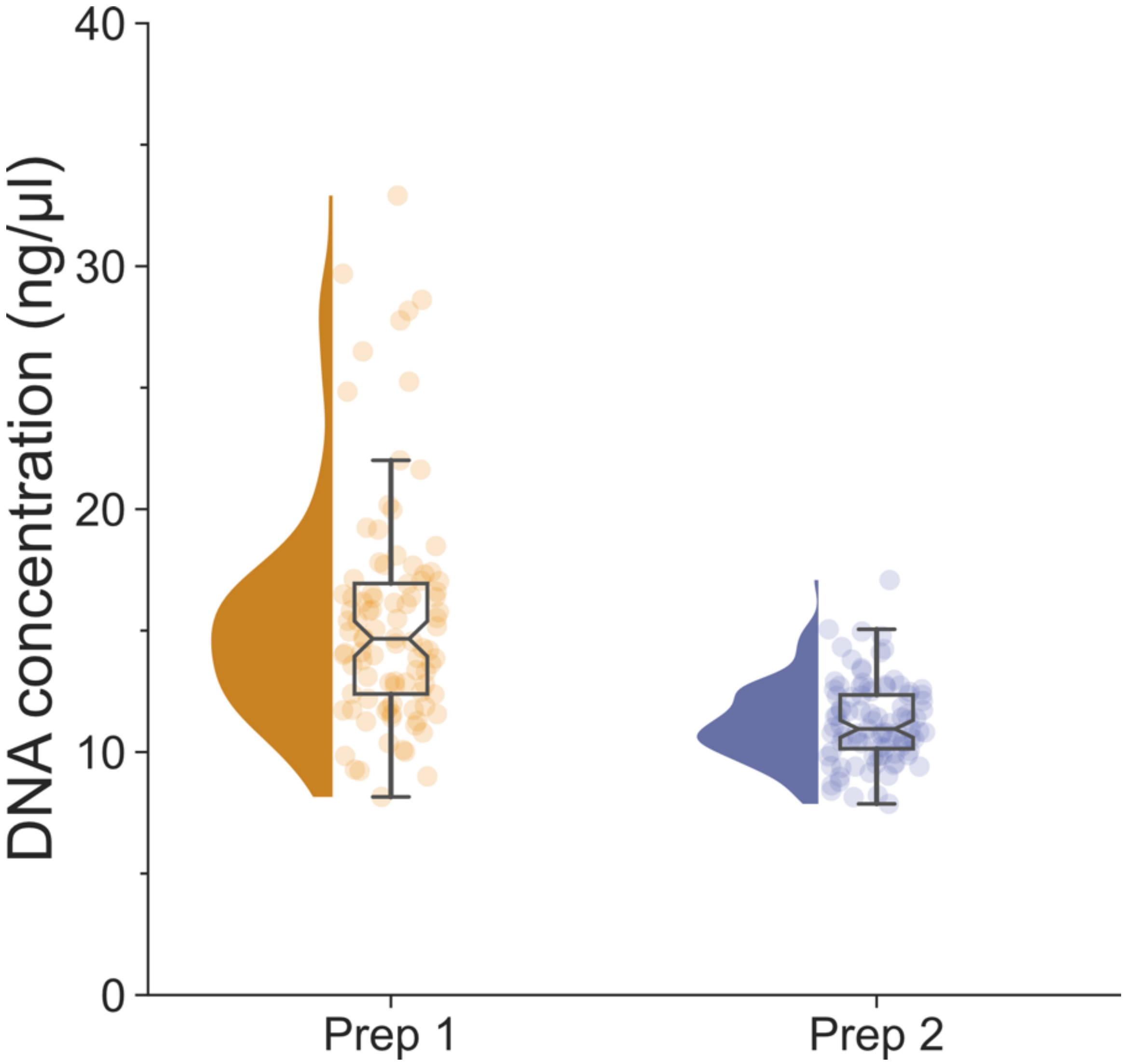

**Supporting Figure 32. The DNA yield of all the samples in Prep 1 and 2 after the SPRI based automated DNA origami purification via liquid handling robot.** The average concentration of the purified DNA origami for plate 1 is at 15.56±0.48 ng/µl. The average concentration of the purified DNA origami for plate 2 is at 11.22±0.17 ng/µl. Errors are the standard error of the means.

### Section 3: Supporting Instruction

This is aimed to provide some short recommendations for adapting the usage of SPRI beads to purify DNA origami.

1. We recommend aliquoting the SPRI beads into tubes, the beads tend to settle down on the bottom of the tube relatively quick, vigorously mixing prior to aliquoting ensure consistent performance of the beads.
2. It is important to mix the beads well prior to use, the beads can be mixed vigorously by hand or by vortex.
3. Due to suppliers could change the SPRI beads components, we recommend that a beads ratio screening should be performed, as the ratio presented here could differ.
4. For different structures, a ratio screen should be performed. In general, we found that ratio between 1.0X to 0.4X works well.
5. We recommend that the volume of the DNA origami should not be less than 20μl. Due to the properties of SPRI beads, the volume can be adjusted to 40μl without affecting the beads performance.
6. After the formation of the bead pellet, during the removal of the supernatant, some beads can be aspirated together with the supernatant, due to the overall yield from this method, it does not impact the overall performance of this method.
7. The last resuspension step, while we resuspended the solution in 40μl to maintain the input and output volume consistency, this can be in any liquid volume, *i.e.* multiple tubes of reaction can be combined into one and elute with smaller volume to concentrate the purified DNA origami.
